# Combined evidence reveals the origin of a rapid range expansion despite retained genetic diversity and a weak founder effect

**DOI:** 10.1101/2025.01.06.631438

**Authors:** Nora M. Bergman, Petteri Lehikoinen, Edward Kluen, Staffan Bensch, Camilla Lo Cascio Sætre, Fabrice Eroukhmanoff, Frode Fossøy, Petr Procházka, William J. Smith, Bård G. Stokke, Craig R. Primmer, Rose Thorogood, Katja Rönkä

**Author notes:** Correspondence should be addressed to Nora M. Bergman.

## Abstract

Many species are currently experiencing range shifts in response to changing environmental conditions, but with potentially serious genetic consequences. Repeated founder events and strong genetic drift are expected to erode genetic variation at the range front, reducing adaptive potential and slowing or even halting the expansion. However, the severity of these consequences for the more common and highly mobile species undergoing environment-driven range shifts (c.f. invasions) is less clear. Here we combined historical observations and contemporary movement data of the common reed warbler (*Acrocephalus scirpaceus*) with genomic evidence from across its European breeding range to (1) infer the origin and (2) quantify the genetic consequences of a recent and rapid northward range expansion. While there were no reductions in levels of nucleotide diversity or allelic richness, nor a signal of founder effect in the directionality index (ψ), our combined dataset approach was able to infer an expansion origin from the southwest. Furthermore, we found that private allelic richness retained a slight but significant linear decline along the colonisation route. These results suggest that high dispersal capabilities can allow even philopatric species to avoid the loss of genetic diversity during rapid range expansions. Nevertheless, if multiple lines of evidence enable identification of an expansion pathway, we may still detect genetic signals of expansion.

## 1. Introduction

Many species across taxonomic groups are undergoing rapid range shifts (Parmesan & Yohe, 2003; Chen et al., 2011) as they track concomitant spatial shifts in suitable environmental conditions (Waldvogel et al., 2020). While moving allows these species to persist or expand in the short term, populations at leading edges are also subject to various evolutionary forces (e.g. strong genetic drift; Excoffier et al., 2009) that affect adaptive potential and thus projections for species’ future resilience (Aguirre-Liguori et al., 2021; Thorogood et al., 2023). However, much of the empirical evidence and development of theory into the genetics of contemporary range expansions (as opposed to longer, post-glacial timescales) has focused on systems involving human-mediated invasions or threatened species. In contrast, much less attention has been paid to environment-driven expansions of geographically widespread species (Garnier & Lewis, 2016; Wallingford et al., 2020), which can have a significant impact on the structure and function of ecosystems (McGeoch & Latombe, 2016). Furthermore, among the studies investigating such environment-driven expansions, most have been conducted with species that remain resident year-round (e.g. wall lizards; Schulte et al., 2013). Migratory species, by comparison, are highly mobile with greater potential for dispersal (Weatherhead & Forbes, 1994) and continued gene flow between the range edge and core areas. Consequently, their range-front populations may show weaker genetic signatures of expansion than what is currently expected, even for rapid expansions (Excoffier et al., 2009; Garnier & Lewis, 2016), but this remains relatively understudied.

In theory, when new areas are colonised, repeated founder effects and small (effective) population sizes are expected to cause strong drift at the expanding range edge (Excoffier et al., 2009; Welles & Dlugosch, 2019). Without sufficient gene flow, genetic diversity is expected to decline and genetic differentiation increase along the expansion axis (Austerlitz et al., 1997; Slatkin & Excoffier, 2012). Before new mutations arise, the new populations can only carry the same or a subset of the alleles present in the source populations. This can lead to clines in allelic and private allelic richness, as it is unlikely that every allele, especially when rare, would be carried along in each founding event or recovered soon through gene flow (Calafell et al., 1998; Jezkova et al., 2015; Szpiech et al., 2008). The effects of drift may have serious consequences for newly founded populations: loss of genetic diversity can cause reduced responses to selection and negative fitness consequences in both theoretical (Peischl et al., 2015) and empirical studies (Hagan et al., 2024; Pujol & Pannell, 2008; Szűcs et al., 2017). In some cases, however, reduced gene flow can also facilitate local adaptation (Kirkpatrick & Barton, 1997; but see Kottler et al., 2021). While there are numerous empirical examples of range expansions showing these expected consequences of drift and repeated founder effects (e.g. Southern flying squirrels *Glaucomys volans*; Garroway et al., 2011, hazel grouse *Tetrastes bonasia*; Rózsa et al., 2016, white-footed mouse *Peromyscus leucopus*; Garcia-Elfring et al., 2017), there are also range expansions where these patterns are weak or non-existent, or have been observed to decrease in severity soon after initial colonisation (Hagen et al., 2015; Robalo et al., 2020). These examples are often of species with high dispersal capabilities (e.g. common crane *Grus grus*; Haase et al., 2019, light-vented bulbul *Pycnonotus sinensis*; Song et al., 2013). Nevertheless, it remains difficult to draw conclusions about fine-scale genomic patterns as many of these studies have relied on low-density markers (mtDNA and microsatellites, but see Adams et al., 2023 for a recent example where high-resolution markers allowed the detection of slight reductions in genetic diversity), or there have been other factors with potential to alleviate founder effects (e.g. relatively small expansion scales or hybridisation with other species). Therefore, it remains unclear how severe and persistent these consequences may be in contemporary range shifts, and particularly for highly mobile species.

Predicting the genetic outcomes of range expansions in species with high dispersal capabilities is further complicated by somewhat contradictory expectations. For example, the retention of genetic diversity is expected to be higher during slow expansions (Excoffier et al., 2009; Garnier & Lewis, 2016), but sufficient gene flow from areas behind the colonisation front can prevent the erosion of genetic diversity even when the expansion is rapid (e.g. invasive starlings *Sturnus vulgaris*, Berthouly-Salazar et al., 2013). Long-distance dispersal (LDD), a context-dependent but characteristically rare and extreme dispersal event (Jordano, 2017), is a key process that can increase gene flow from more distant areas and alleviate initial founder effects. However, very rare LDD events can have the opposite effect and cause greater loss of genetic diversity compared to short-range dispersal, as isolated populations (with low diversity) being founded ahead of the main expansion wave can hinder the settling of later-arriving individuals (Bialozyt et al., 2006). Furthermore, dispersal ability and propensity do not always go hand in hand, with many long-distance migrants moving annually across thousands of kilometres, yet being highly philopatric which can hinder dispersal and gene flow (e.g. Bensch, 1999; Tingley et al., 2012). In general, detecting genetic signatures of range expansions in highly mobile species is likely to require genome-wide datasets that allow for more fine-scaled analyses of patterns that persist along different timescales. For example, some genetic patterns decay more rapidly, such as reduced homozygosity or nucleotide diversity, while others persist for longer, including allele frequency clines or allelic/private allelic richness clines in the direction of the expansion (Allendorf, 1986; Jezkova et al., 2015; Peter & Slatkin, 2013; Szpiech et al., 2008). Comparing these signatures will help to clarify the strength and persistence of founder effects in contemporary range expansions of highly mobile species.

Another problem when investigating the genetic consequences of rapid environment-driven range expansions is that we often lack accurate information on the timing and geographic origin of colonisation (Welles & Dlugosch, 2019). Identifying the source population(s) is vital if we are to make meaningful comparisons of genetic changes during an expansion. However, reconstructing the expansion pathway is a non-trivial task as it requires (a) direct, often anecdotal observations, (b) careful monitoring schemes, or (c) molecular data with sufficient resolution across an appropriate geographical area. An alternative approach is to draw these multiple lines of evidence together to improve confidence in inferring colonisation history and alleviate potential biases associated with any single approach. For example, observational bias can arise with less conspicuous species or those that inhabit hard-to-access habitats (Hughes et al., 2021), and methodological issues remain in commonly used genomic analyses (e.g. Kemppainen et al., 2024). Different lines of evidence allow to elucidate the origins of expanding populations that might have avoided or lost the genomic signals characteristic of a range expansion, as detailed above. Despite these benefits, however, few studies of range expansions take advantage of such a combined-evidence approach, although it is becoming a useful tool in reconstructing e.g. invasion routes (Campbell et al., 2018) or resolving temporal incongruities in species divergence (i.e. ‘total evidence’; e.g. Heritage & Seiffert, 2022).

The common reed warbler (*Acrocephalus scirpaceus*) provides an excellent study system to use combined evidence to detect whether there are genetic consequences in an environment-driven range expansion of a species with high dispersal capabilities. The nominate subspecies (*A. s. scirpaceus*, from here on “reed warbler”) is a long-distant migrant with a large distribution, wintering in sub-Saharan Africa and breeding in reedbeds (stands dominated by *Phragmites*) across Europe, western Turkey, and north-western Africa (Gill et al. 2023). Current knowledge suggests that, at least in the core of their breeding range, reed warblers show a site-fidelity pattern typical of songbirds (Leisler & Schulze-Hagen, 2015): strong breeding philopatry, some natal dispersal, and rarely-detected LDD events (while the definition of LDD is somewhat arbitrary, in Great Britain 96.7% of natal dispersal and 99.9% of breeding dispersal movements of the reed warbler have been <12 km; Davies, 2019). Nevertheless, the species has undergone a rapid expansion at their northern range edge, establishing populations in Sweden, Finland, and Norway as well as all of the Baltic countries during the past ∼150 years. While there are written records of early observations in this region (Leivo, 1937; Sits, 1936; F. E. Stoll as reported in Meder, 1934), the possible range expansion pathways into the north remain ambiguous. Re-encounter data of ringed birds (e.g. Wynn et al., 2022) suggests that reed warblers in Fennoscandia and the Baltic countries migrate to their breeding grounds from the southwest (i.e. through Western Europe), and thus likely indicate that their range expansion occurred along this route. However, it is also possible that colonisation of Finland, in particular, may have been influenced by birds originating from populations more directly to the south or from further to the southeast. Observations of colonisations along these possible routes are available (e.g. Malchevsky & Pukinsky 1983, Glutz von Blotzheim, 1991) but not yet systematically collated. In addition, as reed warblers show a prominent migratory divide in Central Europe (Procházka et al., 2018) and because migration bearing is heritable in birds (Liedvogel et al., 2011), ringing re-encounter data of individuals on migration to and from range edges (e.g. Fransson & Stolt, 2005) could provide a second complementary line of evidence to reconstruct the origins of this species’ range expansion.

Here we combine historical observations, ringing records, and genomic data to address the following two questions: (i) What are the origins of the recent and rapid northward range expansion of the reed warbler in Finland, and (ii) can loss of genetic diversity or clines in allele frequencies be detected along this expansion axis? Genetic differentiation across the reed warbler’s European breeding range is low (Procházka et al., 2011), but a recent study using restriction-site associated DNA sequencing (RAD-seq) data suggests that there should be sufficient structure to detect genetic patterns of range expansion (Sætre et al., 2022, preprint), if such patterns were to persist. Here we complement this dataset with sequencing of additional samples to cover most of the breeding range and all recently colonised northern areas (c.f. Sætre et al., 2022, preprint). We first collate literature of the first observations of breeding in Fennoscandia, northwestern Russia and the Baltic countries in their original languages, and then use ringing records of birds caught during the breeding season in Finland and elsewhere on migration to infer population origins and expansion routes. We then hypothesise that whilst genetic diversity is likely to be mostly maintained along the colonisation route due to gene flow (Procházka et al., 2011; Sætre et al., 2022, preprint), longer-lasting signals of past founder effects in allele frequency clines or patterns in (private) allelic richness may still persist. This approach allows us to inspect the fine-scale genetic signals of past founder effects in a documented, environment-driven range expansion, providing a novel example for understanding the conditions under which species can successfully respond to ongoing environmental change.

## 2. Materials and Methods

### 2.1 Literature search for historical observations

We conducted a literature search to locate as many historical records of local first occurrences of reed warblers and their distributions in Northern Europe (e.g. Denmark, Estonia, Finland, Latvia, Lithuania, Norway, Russia, Sweden) as possible. We relied mostly on published observations, supplemented by secondary sources such as yearbooks of local birding societies and handbooks. To find records from the areas of the former Soviet Union and East Prussia, we contacted a local expert (N. Chernetsov) for guidance.

### 2.2 Migratory direction

We collated all available data from reed warblers ringed in Finland and re-encountered outside of Finland (years 1969-2023, n = 326) from the databank of the Ringing Centre at the Finnish Museum of Natural History. As Finland is at the species’ range edge and there should be no individuals migrating through the country to other areas, we used ringing data from all months. However, these should be interpreted as ringing locations instead of confirmed natal or breeding sites, as migration continues during much of the breeding season. To investigate any possible contributions from different sides of the species’ migratory divide, we split the Finnish data into a western (n = 173) and an eastern group (n = 153) arbitrarily by the mean longitude of the site of ringing to measure differences in mean migratory direction and individual migration routes. Mean individual migratory directions were calculated as the consistent compass bearing between the ringing and re-encounter events. The mean direction of each group was calculated based on re-encounters south of 49°N (western group n = 26, eastern group: n = 33) to avoid the large number of birds controlled in Belgium (due to high catching effort) affecting the mean (see Fransson & Stolt, 2005). Furthermore, this allowed us to avoid bias from re-encounters made near the breeding area, since directions calculated based on short distances may reflect shorter-scale movements (e.g. post-fledging and post-breeding movements; Chernetsov, 1998; or avoiding geographic barriers such as the Baltic Sea) and less reliably represent innate migratory direction (e.g., Helbig, 1992). Finally, to statistically test whether ringing longitude affects migratory direction, we fit the following simple linear models in R (v4.3.0; R Core Team 2023) to the dataset containing re-encounters made south of 49°N: 1) recovery longitude ∼ ringing longitude, 2) recovery direction ∼ ringing longitude. Residual diagnostics were assessed using the DHARMa package (Hartig, 2024) and showed no significant deviations from model assumptions.

### 2.3 Population genomics

#### Sample collection, DNA extraction, and RAD-seq library preparation

We collected or received blood samples from reed warblers representing 20 sites at least 50 km apart from each other across the species’ European breeding range (Fig. 1). A total of 171 individuals (n = 5–10 per sampling site) were included in the final dataset. Blood samples were taken from resident birds during the breeding season in years 2002-2019 and stored in either Queen’s lysis buffer (+4°C or −20°C; Seutin et al., 1991), ethanol (+4°C) or on FTA cards^TM^ (Whatman, Maidstone, United Kingdom) (room temperature). For sampling details, see Table S1.

**Figure 1.**
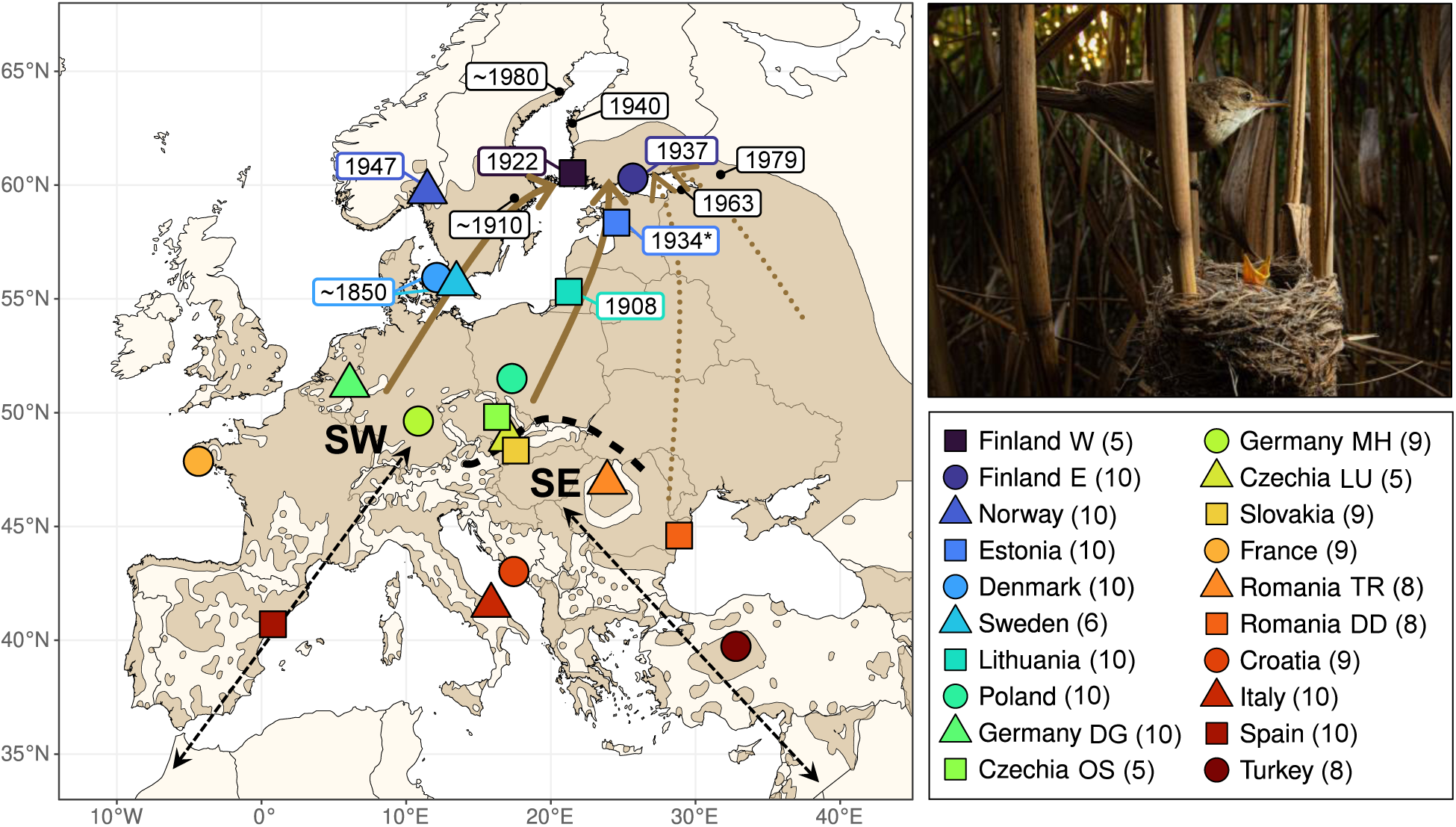
Left: Map of the reed warbler’s current breeding range (in brown) in the study area (distribution data from BirdLife International and Handbook of the Birds of the World, 2024). The hypothesised expansion pathways to Finland are illustrated with brown arrows, of which solid arrows denote the proposed routes based on the results of this study. The species’ migratory divide in Central Europe is indicated with a black dashed line (following Procházka et al., 2013), with populations on the different sides of the divide using either a southwestern (SW) or a southeastern (SE) route between their wintering and breeding grounds (general directionality of these routes indicated with black dashed arrows). The sampling sites for molecular data are marked on the map with filled shapes coloured by latitude order, legend (bottom right) shows the site names with the number of individuals used in molecular analyses. The year of first recorded occurrence is denoted on the map for respective sampling sites or other areas (small black dots) within the newly colonised range. The * by the Estonian first occurrence indicates the year of the first record of the species in over 60 years, although a single nest was reported already in 1870. Top right: A common reed warbler at its nest. Photograph: Deryk Tolman.

Genomic DNA was extracted using a QIAGEN DNeasy Blood & Tissue Kit (Qiagen Inc., California, U.S.A) at the University of Oslo or the University of Helsinki. The samples were first handled according to their storage medium, after which the manufacturer’s protocol was followed with minor changes (Supplement S2). After extraction, the purity and DNA concentration of the eluates was assessed using a NanoDrop 2000 spectrophotometer (Thermo Scientific) and a Qubit 4 fluorometer (Invitrogen). The DNA concentration of each sample was normalised to 20 ng/μl before sequencing. Gel electrophoresis was used to ensure that the DNA fragments were of desired size (molecular weight > 10 kb).

Single-digest, single end RAD-seq libraries were prepared at Floragenex, Inc. (Oregon, U.S.A) for a total 285 samples (including 12 replicate controls, as well as samples sequenced as a part of another project and not analysed further here), following Baird et al. (2008). DNA was digested with the *Sbf*I enzyme and sequenced on an Illumina HiSeq 2000 platform. Each sample was barcoded and sequenced on two separate lanes. Samples were sequenced on three plates, one in 2017 (read length 91bp) and two in 2021 (read length 150bp). Replicate samples were included between all plates to allow the detection of potential batch effects, and samples sequenced in 2021 were randomised across the two batches to decouple the biological signal from any batch differences (Meirmans, 2015).

#### Sequence processing and reference alignment

The raw sequence quality was assessed using FastQC (Andrews, 2019), after which the data was cleaned and processed with STACKS (v2.62; Catchen et al., 2013; Rochette et al., 2019). The raw reads from each sequencing lane were cleaned and demultiplexed separately with the STACKS *process_radtags* program, using default settings: allowing for maximum one mismatch in the barcode or restriction site, and discarding the read if the average Phred score in a sliding window drops below 10. This resulted in a total of 2.14 billion reads across all three sequencing batches, out of which 90.6% were retained. The non-retained reads were dropped due to ambiguity in barcodes (8.6%) or restriction sites (0.2%), or a low quality score (0.6%). All reads were truncated to equal length (91bp), and sequences from the two lanes of each plate were combined to increase total coverage. The cleaned reads were aligned to the *A. scirpaceus* reference genome (Sætre et al., 2021) using BWA-MEM (v0.7.17-r1188; Li, 2013) with default settings. The resulting BAM files were sorted using SAMtools (Li et al., 2009), and the STACKS *ref_map.pl* program was run to assemble the loci according to alignment positions, and to perform SNP calling in each sample. The program retained 69.6% of the primary alignments, the rest were dropped due to insufficient mapping qualities (25.7%), excessive soft-clipping (1.4%), or as unmapped (3.3%). From these, 452 673 RAD loci (containing 997 942 SNPs) were built with an effective per-sample mean coverage of 47.1x (SD=16.8x, min=15.1x, max=111.7x).

To check for any batch effects between our three sequencing plates (2017, 2021_plate1, and 2021_plate2), we ran STACKS *populations* without filters for the 12 control samples that were re-sequenced across plates. We checked the relatedness of each replicate pair (VCFtools --relatedness; v0.1.16; Danecek et al., 2011), which corresponded to the same individual in all samples except one, where a similar relatedness value to another sample indicated likely mislabelling. These samples were excluded from further analyses. We then checked that the control samples cluster by sample identity and not by batch in a principal component analysis (O’Leary et al., 2018), which showed no signs of batch-based clustering across any of the three batches (Figure S3). With VCFtools, we output the observed and expected heterozygosity, mean depth, and missing data per individual and compared sample values across batches using paired, two-tailed t-tests. The two plates sequenced in 2021 did not differ significantly from each other in any of the tested values (Table S4, Figure S5), and therefore the 2021 controls were treated as one group in the comparisons with the 2017 plate. Between the 2021 and 2017 batches, we detected small but statistically significant differences (p < 0.05) in all tested statistics (Table S4, Figure S6), observed heterozygosity being on average 3.7% lower and expected heterozygosity 0.1% lower in the 2017 batch. As even slight differences in heterozygosity due to technical artefacts might produce a false signal when biological differences are small, we only used samples sequenced in 2021 (i.e. Datasets 3 & 4, Table S7) for the most sensitive analyses regarding allele frequency clines and genetic diversity.

#### Creating the filtered datasets for population genomic analyses

We first ran STACKS *populations* without filters and calculated the mean sequencing depth and missingness per site and individual using VCFtools; no individuals were discarded based on these. With VCFtools, we extracted a list of loci containing sites with minimum mean depth 10 and maximum mean depth corresponding to 2x mean depth of all sites. These loci, together with all loci on sex chromosomes, were converted into a blacklist for *populations*. After re-running *populations* with this blacklist and basic quality filters, we ran an exploratory PCA using the glPca function in the R package *adegenet* (Jombart & Ahmed, 2011) and excluded two extreme outlier individuals from further analyses (likely misidentified species or subspecies, see Arbabi et al., 2014). We then ran *populations* without these individuals but with otherwise same filtering settings as in the previous step and calculated the relatedness between samples (VCFtools --relatedness). Only one individual from each pair or group of close relatives (1^st^-3^rd^-degree) was kept (e.g., Wang, 2018). We also removed two samples with a pairwise relatedness value corresponding to a duplicate or monozygotic twin, likely indicating contamination or mislabelling. STACKS *populations* was run again without control samples and mean depth was re-calculated for a new blacklist, again containing the sex chromosomes and sites with mean depth <10 or >2x the mean depth of all sites. To determine a thinning threshold for analyses that assume unlinked markers, we used the PopLDdecay software (v3.42; Zhang et al., 2019) to calculate the decay of linkage disequilibrium over physical distance.

Finally, we downsampled the data to a maximum of 10 individuals (n = 5–10) per sampling site to avoid the effects of uneven sampling on the genomic analyses (e.g., Lawson et al., 2018). STACKS *populations* was run to create the final datasets shown in Table S7. In all four datasets, we required a locus to be present in at least 80% of individuals (-r) in 75% of the sampling sites (-p) to reduce missing data, applying these filters haplotype-wise (-H) in the datasets with full RAD loci. We set a minimum minor allele count threshold to exclude singletons and private doubletons (--min-mac 3; Linck & Battey, 2019), a maximum heterozygosity threshold of 70% across all samples (--max-obs-het 0.70), and applied the blacklist described above (excluding sex chromosomes to prevent individuals clustering by sex instead of population structure, as well as low- and high-depth loci to prune out potential false homozygotes and paralogs). To exclude any remaining genotyping errors, we scanned for markers out of Hardy-Weinberg equilibrium in all populations with n = 10 (to increase statistical power), but there were none left to be removed at this stage. For the analyses assuming independent SNPs or RAD haplotypes (Table S7), we used VCFtools to thin the data according to the calculated LD decay distance (Figure S8) and ran STACKS *populations* a final time using just a whitelist of the kept markers: SNPs min 3000 bp apart (**Dataset 1)**, or RAD loci min 3000 bp apart (**Dataset 2**). To be certain that the slight batch effect does not drive the results even in the most sensitive analyses, we additionally created two datasets with only the batches sequenced in 2021: SNPs min 3000 bp apart (**Dataset 3**) and a non-linkage-pruned dataset containing all SNPs within the RAD loci (**Dataset 4**).

#### Population structure and differentiation analyses

While population structure analyses are not explicit inferences of colonisation history, they provide useful information about patterns of genetic variation and connectivity between populations. As recommended for population structure inference (Linck & Battey, 2019), we used complementary non-parametric and model-based methods to depict the structure and differentiation in our data. The specific datasets used in each analysis are listed in Table S7.

We performed a principal component analysis (PCA) for the full, linkage-pruned SNP dataset (Dataset 1) using the glPca function in the R package *adegenet* (Jombart & Ahmed, 2011) and plotted the results with *ggplot2* (Wickham 2016). To match the PCA orientation with map orientation for more intuitive viewing, the PC axes were inverted by multiplying PCA scores by −1, a visual adjustment that does not affect the relationships between samples. Correlations between the PCs and geography were also calculated using the inverted PCA scores. To use the resolution provided by full RAD haplotypes for analysing population structure, we ran the population inference package fineRADstructure (v0.3.2; Malinsky et al., 2018) for the full, linkage-pruned RAD locus dataset (Dataset 2), allowing maximum 10 SNPs per locus (-n 10). fineRADstructure’s MCMC clustering algorithm was run with 100,000 burn-in iterations, followed by 100,000 iterations sampled every 1000 iteration steps (-x 100000, -y 100000, -z 1000), and tree-building algorithm with 10,000 iterations (-m T, -x 10000). To disentangle any patterns of discrete structuring that cannot be explained by isolation by distance only, we used the spatial, model-based clustering package conStruct (v1.0.5; Bradburd et al., 2018) for the full, linkage-pruned SNP dataset (Dataset 1). We first ran a cross-validation procedure implemented in conStruct to select between spatial and non-spatial models and the number of discrete layers (values of K). Cross-validation was run for K = 1–5, with four repetitions with each value of K, 5000 iterations per repetition, and a training proportion of 0.9. To choose the most informative value of K for our data, we required that each new layer should contribute at least 5% to the overall covariance of the model. The diagnostic plots for MCMC performance were checked for all repetitions of the chosen value of K to confirm that the chains were well-mixed. To visualise the conStruct results on map, we calculated the average ancestry proportion of each layer across the samples from each sampling site. Finally, we estimated the relative genetic differentiation between sampling sites. Pairwise F_ST_ values and their statistical significance, based on 5000 bootstrap iterations, were calculated using the R package StAMPP (v1.6.3; Pembleton et al., 2013) for the full, linkage-pruned SNP dataset (Dataset 1).

#### Origins of the range expansion

To test whether a genetic pattern of past founder effects persists, we used an approach by Peter & Slatkin (2013, 2015), implemented in the RangeExpansion R package. The directionality index (ψ) detects allele frequency asymmetries, caused by past founder effects, between pairs of populations. Clinal variation in ψ is still credited as the most sensitive statistic for detecting signals of range expansions in genomic data, its power decaying slower than that of F_ST_ or heterozygosity (Kemppainen et al., 2024; Peter & Slatkin, 2013). We calculated the pairwise ψ values for all sampling sites in the batch-effect-free, linkage-pruned SNP dataset (Dataset 3) with the modified function (get.all.psi.mc.bin) from Kemppainen et al. (2024), which fixes a major bug in the original package by inverting the polarity of the result matrix and applies a binomial test to calculate the significance of each value. To further distinguish from false positives potentially caused by range boundary effects, we rescaled the mean absolute value of ψ by the mean pairwise F_ST_ in this same dataset (ε = |ψ| / F_ST_) and calculated the value of τ (the strongest positive r^2^ value between any population pair from the non-TDoA method; Kemppainen et al., 2024; Ramachandran et al., 2005). We then compared these values to the simulated data in Kemppainen et al. (2024) to see whether the signal in our data falls within the prediction interval for a signal of real range expansion at any significance level.

#### Genetic diversity

To compare levels of genetic diversity across the sampled range of the reed warbler, we used STACKS *populations* to calculate nucleotide diversity (π) for each sampling site using the batch-effect-free, non-linkage-pruned full RAD locus dataset (Dataset 4), including both variant and invariant sites (e.g. Korunes & Samuk, 2021). Allelic richness (AR) and private allelic richness (PAR) were calculated using ADZE (v1.0; Szpiech et al., 2008) for the same dataset (Dataset 4), allowing a maximum of 10% missing data per locus. The private allele calculations were performed for a subset of sampling sites in this dataset (excluding Spain and Romania to only quantify patterns in the recently colonised range) to detect possible losses of rare alleles during the range expansion. For both AR and PAR, the number of samples per locality was standardised to the smallest number of allele copies across all sampling sites (G = 10, i.e. five diploid individuals) as uneven sample sizes impact the number of distinct alleles observed (Kalinowski, 2004; Szpiech et al., 2008). We tested the correlation of each statistic with latitude using a simple linear regression. For π and AR, we also compared the mean values between the recently colonised range (Fennoscandia and the Baltic countries) and the rest of the sampled range using two-tailed t-tests. As PAR can be affected by the sampling scheme (i.e. if sampling sites and therefore local populations are closely related and therefore share the same alleles, few alleles will be private to any single location), we repeated the linear regression against latitude excluding i) one of the Finnish sites FIE due to its geographical proximity to FIW, and ii) both FIE and DK due the proximity of the latter with SE. Additionally, we tested the correlation between PAR and latitude along possible expansion routes to Finland along iii) the western coast and iv) the eastern coast of the Baltic Sea, using unique sampling sites in each model (Figure S15).

## 3. Results

### 3.1 Historical observations suggest a southwestern expansion origin

During the 19^th^ and early 20^th^ century, reed warblers were documented expanding northwards along both sides of the Baltic Sea. In terms of latitude, the colonisation wave along the western coast through Scandinavia seems to have been a few decades ahead of the expansion front advancing along the eastern coast of the Baltic Sea. In the mid-1800s (the beginning of available observation data for the species), reed warblers were likely breeding in most of mainland Denmark (Dybbro, 1976; Løppenthin, 1967), the regions south of Kaliningrad, Russia (at the time Königsberg, East Prussia; Tischler, 1914), and a few locations in southern Sweden (Risberg, 1990). The species expanded rapidly from the late 1800s onwards, in the west reaching the Stockholm area around the 1910s (Risberg, 1990; Svensson et al., 1999), and in the east settling in northern parts of the Curonian Lagoon in the early 1900s (Tischler, 1941). The expansion reached southwestern Finland in the 1920s (Turku 1922, Åland islands 1926) (Leivo, 1937; Wikström, 1945). The colonisation wave through the Baltic countries followed closely: reed warblers were first found breeding in both Latvia and Estonia during the early 1930s (Riga 1931, Pärnu and Matsalu Bay 1934, although a single, possible nesting has been reported from Estonia already in 1870) (Mang, 1936; Sits, 1936; Tischler, 1941; Russow, 1874 as cited in Sits, 1936). However, it is possible that the first colonisers went undetected, as multiple individuals and nests were found in all these locations already in the year of the first observation. The first finding from the Helsinki region in Finland was made in 1933 but based on how common and widespread reed warblers were found to be in 1937, Leivo (1937) suggested that the species might have already occurred in the area some years before. It therefore seems possible that the two expansion waves, from the southwest (Sweden) and south (Estonia), both reached Finland in the 1920s-30s. In Norway, the first confirmed breeding was in 1947 at Lake Borrevatn near Oslo, although the first sighting was made a decade earlier on the island of Utsira at the SW coast of the country (Haftorn, 1971; Løvenskiold, 1947). To the easternmost tip of the Baltic Sea in the Saint Petersburg region, Russia (at the time Leningrad, USSR), reed warblers only arrived in the 1960s, 30-40 years after the first records from neighbouring areas in Finland and Estonia (Malchevsky & Pukinsky, 1983). A distribution map from the 1950s shows that the species was absent directly east of the Leningrad area 10 years before the first observation near Saint Petersburg, and ∼500 km away in the south (Dement’ev et al., 1954/1968). Observations support that the expansion proceeded from west to east and from coastal areas inland (Malchevsky & Pukinsky, 1983), making east an unlikely origin for the Fennoscandian and Baltic reed warbler range expansion.

### 3.2 Migratory direction supports expansion from the SW side of the migratory divide

As expected under the hypothesis that the reed warbler’s range expansion originated from the southwestern side of the species’ migratory divide, none of the individuals ringed in Finland (n = 326) were re-encountered along the southeastern migration route. However, there was some variation in where birds were re-encountered along the SW migration route. For recoveries south of 49°N (n = 59), the birds ringed in eastern parts of Finland had a marginally significant tendency to be recovered at more eastern longitudes, and vice versa (β = 0.98, SE = 0.51, t = 1.92, p = 0.059), but the correlation between ringing longitude and recovery direction was not significant (β = 0.50, SE = 0.72, t = 0.695, p = 0.49). Interestingly, birds ringed in the western (n = 173) and eastern (n = 153) parts of Finland were both found to be migrating to and from their breeding grounds along either side of the Baltic Sea. While most individuals seem to cross the Mediterranean Sea through the Iberian Peninsula, the re-encounter data suggest that Finnish reed warblers may rarely use the Italian peninsula for crossing (Fig. 2), during both the spring and autumn migrations (Figure S9).

**Figure 2.**
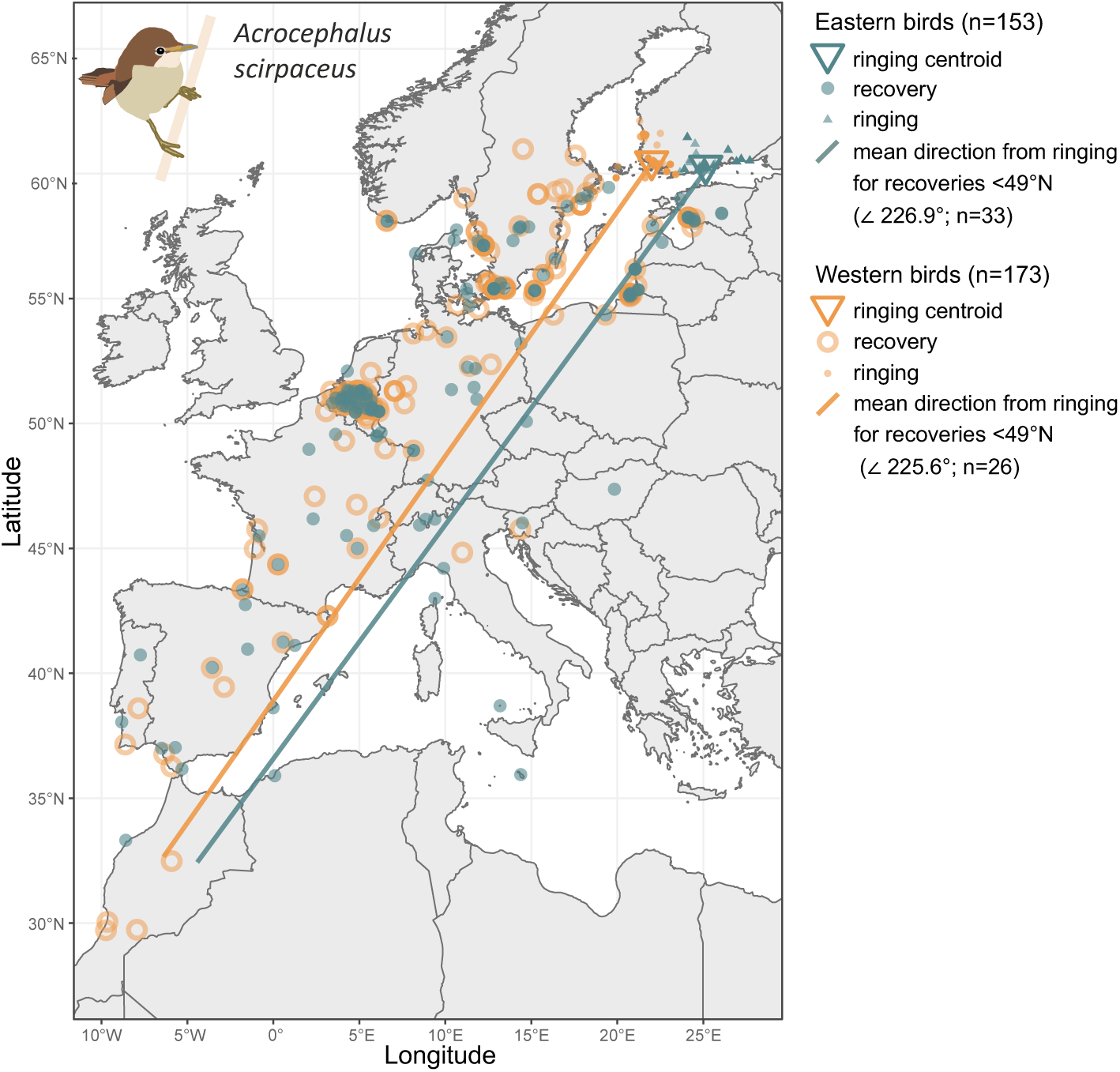
Map of all re-encounters from reed warblers ringed in Finland. The re-encounters are divided into a western (orange) and an eastern (teal) group based on the mean longitude of the ringing site. Opacity indicates multiple, overlapping points. Mean migratory direction for each group is calculated from re-encounters made south of 49°N. For individual trajectories during spring and autumn migrations separately, see Figure S9.

### 3.3 Weak but significant geographically continuous population structure

The principal component analysis indicated that the overall population structure of the reed warbler largely reflects geography across its European breeding range. PC1 (1.13% of variation) aligned samples along a south-north cline (Pearson correlation with latitude: r^2^ = 0.887, 95% CI = 0.922 - 0.956, p < 0.001), PC2 (0.82%) reflected longitude (r^2^ = 0.467, 95% CI = 0.595 - 0.756, p < 0.001), and PC3 (0.76%) captured differentiation among the southernmost sampling sites, separating most Spanish individuals and one Italian bird from the rest of the samples (Fig. 3A-B). However, PC1 also positioned the most recent northern expansion in Norway at the end of the genetic continuum, despite samples being collected 1 degree south of Finland. The results from fineRADstructure concurred with the PCA: individuals were generally clustered by sampling site and geography in the coancestry matrix, but as many of the main splits were not well-supported as separate clusters (posterior probability < 95%) this indicated continuous rather than discrete structuring, especially in Central and Northern Europe (Fig. 3C). The same patterns were reflected in F_ST_, where in general, differentiation increased with increasing geographic distance (Fig. S10). Pairwise F_ST_ values were low, but mostly statistically significant (p < 0.05); of these significant comparisons, pairwise F_ST_ ranged from 0.021 between eastern Finland and Turkey to 0.001 between e.g. eastern Finland and Norway.

**Figure 3.**
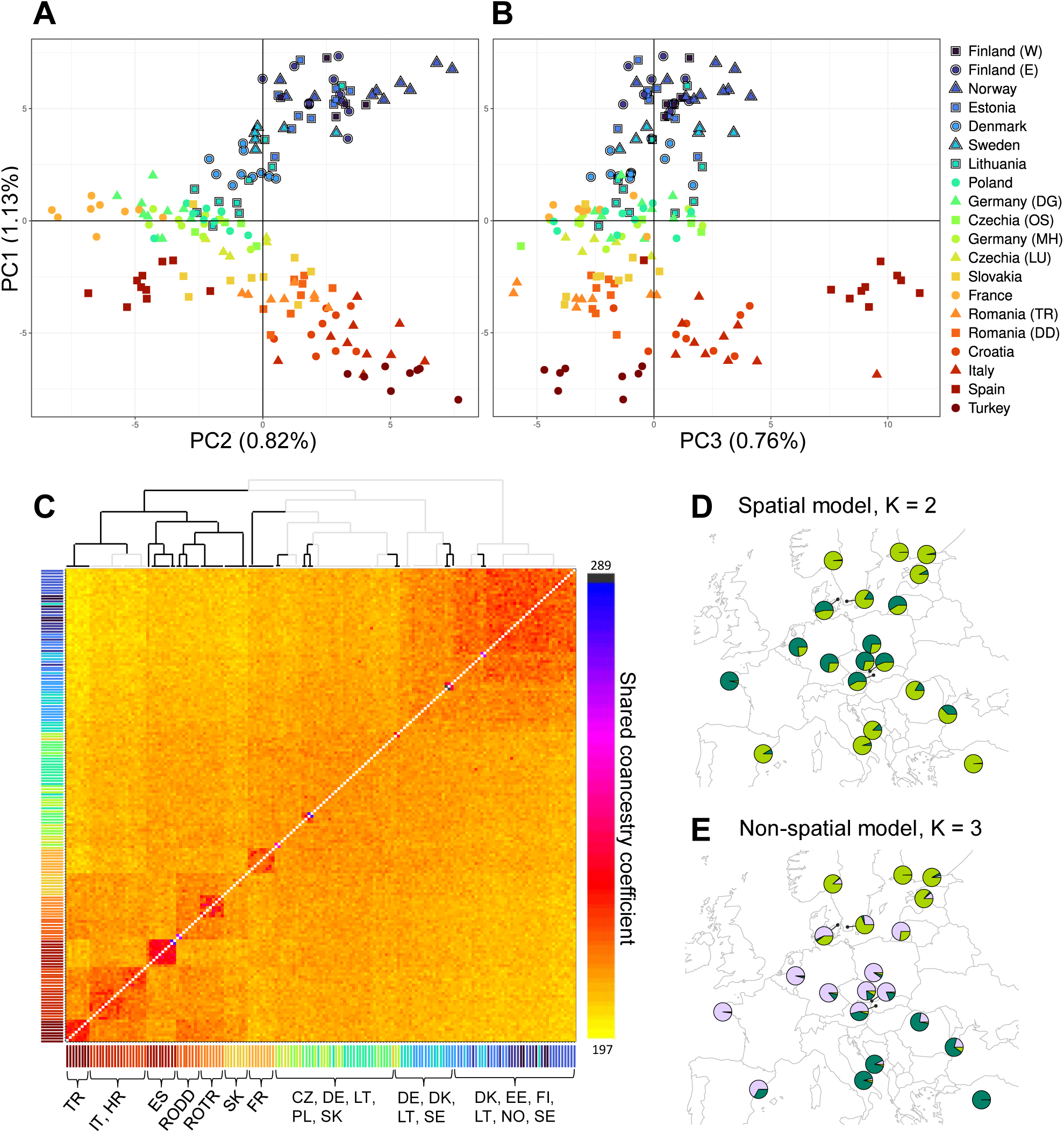
Population structure across the reed warbler’s European breeding range. (A) The two first principal components of the PCA show geographically continuous structure. As a visual adjustment, the PCs have been rotated to match the orientation of the sampling map. Each data point represents an individual, the colour and shape correspond to its sampling site (legend next to panel B). Samples outlined in black are part of the recently colonised range edge with historical observation data available. (B) PC1 against PC3, which captures genetic structuring among the southernmost sites. (C) The fineRADstructure coancestry matrix also shows clustering by geographical proximity (colours of individuals match legend of A & B) with more pronounced population structure in the southern parts of the sampled range. The branches of the tree with high support (> 95% posterior probability) are shown in black. Each row and column correspond to an individual and its genetic relatedness to every other individual in the data, with heatmap colour corresponding to relative genetic similarity. (D) The overall best model in the conStruct analysis, a spatial model with K = 2, suggests the presence of two genetic clusters after accounting for isolation by distance. However, there is very little genetic variation separating the two layers and all replicates did not converge on the same split. The pie charts represent the average proportion of ancestry drawn from each layer K in each sampling site. (E) The best non-spatial conStruct model (K = 3) shows geographical clustering in line with the other non-spatial analyses.

The spatial population analysis conStruct showed that population structure could be largely explained by isolation by distance (IBD), and any additional, discrete patterns were only weak and received mixed support. Using the package’s cross-validation procedure, a spatial model had a higher predictive accuracy than the corresponding non-spatial model across all tested values of discrete layers (K) (Figure S11A-C). As adding a third layer (K) contributed < 5% to the total covariance (Figure S11D), the spatial model with K = 2 was considered the most relevant description of the data (Figure 3D). However, the amount of variation separating the two layers in this model was so small that replicates did not converge on the same split of layers, despite good MCMC diagnostics. In Figure 3D, the most common split for the K = 2 spatial model is shown (2 out of 4 replicates converged). This split indicated that the northern and southern sites would be more similar to each other than expected under IBD alone, but this result should be interpreted with caution as the discrepancies between replicates and the small amount of variation separating the two layers in the K = 2 model (Figure S12) suggest that a spatial model with K = 1 might be the most biologically relevant model (i.e. nearly all spatial variation could be explained by IBD). The non-spatial model (comparable to an ADMIXTURE model; Alexander et al., 2009; Bradburd et al., 2018) that best described the data was K = 3, which again showed geographical structuring in line with the results from all other non-spatial structure analyses (Fig. 3E).

### 3.4 No detectable signal of range expansion in the directionality index

No allele frequency clines were detectable with sufficient confidence to distinguish from an equilibrium scenario. Directionality index (ψ), quantifying directional change in allele frequencies across the sampled range, was very low in all pairwise comparisons between sampling sites (absolute values of |ψ| = 0.0006**–** 0.021). The proportion of significant comparisons (1/66, 1.5%) was lower than the proportion of false positives expected under the null hypothesis of no expansion (5%, Kemppainen et al., 2024). The values of |ψ| fall clearly below the cutoff |ψ| > 3 used by Peter & Slatkin (2013, 2015). With the normalised value of ψ (ɛ = |ψ| / F_ST_ = 0.92) and the highest pairwise r^2^ from non-TDoA (τ = 0.29), our data do not cross the lower limit of the prediction intervals of a real signal of range expansion at any significance level, based on the simulations in Kemppainen et al. (2024).

### 3.5 Genetic diversity

Levels of nucleotide diversity (π) and allelic richness (AR) were highly consistent across the range and showed no reductions in the newly colonised sites (Fig. 4A), but there was a significant reduction in private allelic richness (PAR) with increasing latitude in the recently colonised range (Fig. 4B). The mean value of π ranged from 0.00586 (Sweden) to 0.00608 (Poland) per sampling site, with no linear correlation with latitude or significant difference in mean π between recently colonised sites (Fennoscandia and the Baltic countries) and the other sites (Supplement S13). Values of AR ranged from 1.469 (Spain) to 1.481 (Poland), also with no latitudinal correlation or mean difference between the recently colonised and other sites (Supplement S13), even though allelic richness should retain the signal of founder effects longer than frequency-based measures of genetic diversity, such as π (e.g. Greenbaum et al., 2014). However, private allelic richness (PAR) in the recently colonised areas and the nearest range core sites (Fennoscandia, Baltic countries, Germany, and Poland) showed a highly significant negative relationship with latitude (r^2^ = 0.92, β = −3.96 × 10⁻^4^, SE = 4.31 × 10⁻^5^, t = −9.19, p = 3.73 × 10⁻^5^). PAR was highest at the range core and decreased linearly towards the north, as would be expected under a scenario of sequential founder effects. The difference between the highest and lowest PAR value (Poland: 0.01703, Finland W: 0.01279) corresponds to a difference of 778 alleles in the expected number of private alleles at the standardised sample size used (G = 10, for estimates with different values of G, see Figure S14). The statistically significant pattern persisted even when excluding sites in close geographical proximity (p = 0.0004 when excluding Finland E, and p = 0.002 when excluding Finland E & Denmark; Figure S15A-B). There were similar, linearly declining trends in PAR also when looking at the expansion routes along both sides of the Baltic Sea separately, although the relationship only reached significance along the eastern coast (p = 0.02; Figure S15C-D).

**Figure 4.**
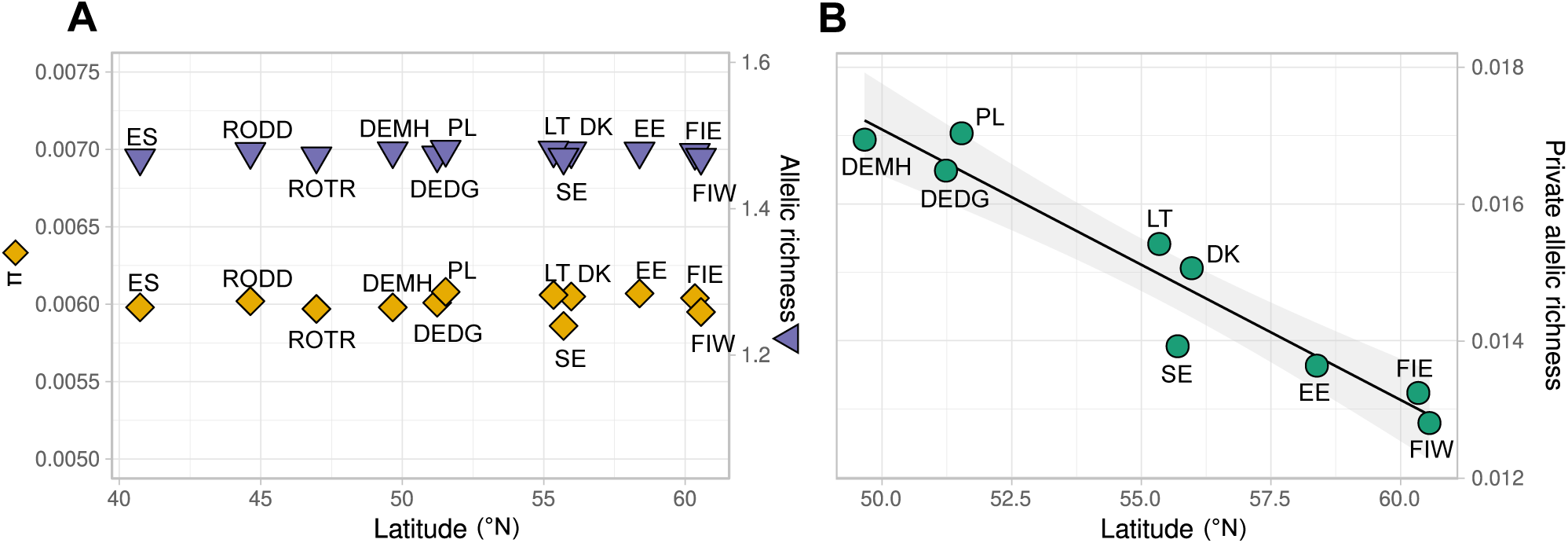
(A) Levels of nucleotide diversity π (orange diamonds) and allelic richness (purple triangles) are highly similar across the sampled range and show no pattern over latitude. (B) Private allelic richness decreases significantly with latitude within the area of interest for the recent, northward expansion. For private allele calculations across the whole data set (including Spain and Romania), see Figure S16.

## 4. Discussion

In a world characterised by rapid environmental changes, it is important to understand the origins and consequences of recent environment-driven range shifts, as the loss of genomic variation following an expansion can impact the population’s resilience in the face of further environmental challenges (Kardos et al., 2021; Thorogood et al., 2023). Here we provide an example of a weak but persistent genetic signal of founder effects despite the absence of losses in genetic diversity after a rapid and extensive environment-driven range expansion of a migratory passerine, the common reed warbler. Despite the expansion spanning more than 1000 km in the past 100-150 years, we did not detect any losses in nucleotide diversity (π) or allelic richness in the newly colonised areas at the northern range edge, nor any signal of founder effects in the directionality index (ψ). However, there was a slight but significant linear decline of private allelic richness in the direction of the expansion. The European breeding range of the reed warbler was characterised by weak, continuous population structure mainly reflecting geographic distance, although we discuss the possibility that accelerated differentiation during repeated founder effects could lead to a similar pattern. The genomic patterns were consistent with the expansion origin estimated from historical observations and migratory direction, all supporting a southwestern origin for the species’ northward expansion to Finland through both Fennoscandia and the Baltic countries.

### 4.1 Complementary evidence for southwestern origin of expansion

Even though colonisation histories can be complex and resolving all details difficult if not impossible, combining independent lines of ecological and genomic evidence allowed us to reconstruct a consistent and well-supported picture of the expansion routes and its associated genomic consequences. While the historical observations provided valuable reference points and suggested an expansion proceeding from the southwest, observational data of species’ range expansions is often incomplete or inaccurate due to e.g. spatial bias in monitoring effort (e.g. Hughes et al., 2021) and does not alone provide means to directly assess the origin of the colonising individuals. However, ringing data also supported an expansion origin from the SW side of the species’ migratory divide in Central Europe, as none of the reed warblers ringed in Finland were recaptured along the SE route (whilst a few individuals were found to cross areas where local breeding birds are SE-migrating, e.g. Hungary or Italy, their migratory direction was still southwestern). The linear decline in private allelic richness provided additional evidence for an expansion pathway from the south and southwest. The pattern is not compatible with a scenario in which there would be a major contribution from Southeastern Europe (exhibiting comparatively high PAR; Fig. S16) or an unsampled population, as this should increase private alleles at the range edge (e.g. Garcia-Elfring et al., 2017). The cline also indicates genetic mixing between the expansion waves advancing on different sides of the Baltic Sea, as Finland should show a higher number of private alleles than the other sites along either side of the sea if it was the meeting point of two isolated expansion waves, each with their own set of stochastically retained alleles. An alternative explanation could be that only the expansion from one side of the sea reached Finland, but the historical observations as well as the lack of population structure between Scandinavia and the Baltic countries suggest otherwise. Furthermore, the continuous pattern of genetic differentiation connecting sampling sites in Central and Western Europe to the recently colonised range points towards shared ancestry (Estoup & Guillemaud, 2010). While relying only on genomic data can introduce a source of bias if some potential source populations are not sampled (Estoup & Guillemaud, 2010), this is unlikely in the case of our study. Although we were not able to include samples from more eastern sites (e.g. Russia), the historical observations were detailed from the time when the expansion proceeded into the St Petersburg region from the 1960s onwards, documenting a chronologically advancing expansion from southwest along the southern coast of the Gulf of Finland (Malchevsky & Pukinsky, 1983).

### 4.2 Why was overall genetic diversity not affected by the range expansion?

Despite the extensive colonisation distance and fast pace, the overall genetic diversity showed no reductions at the range front. The signal of past founder effects was only retained in the rarest alleles in the data, i.e. those that were private to only a single sampling site. Even the directionality index, which has until recently been considered as the most sensitive statistic to range expansions (Kemppainen et al., 2024), and allelic richness, which often also retains a pattern when private allelic richness does (e.g. Heppenheimer et al. 2018; Swaegers et al., 2015, but see Jezkova et al., 2015 for a case where only reductions in private allelic variation persisted after a postglacial expansion of the Merriam’s kangaroo rat *Dipodomys merriami*), showed no signal of expansion. Most previous studies report at least slight reductions in genetic diversity following expansions of species with high dispersal capabilities (Adams et al., 2023; Berthouly-Salazar et al., 2013; Pierce et al., 2014), and there are only a few earlier cases reporting no losses of diversity following expansions of similar scale, without contribution from factors such as hybridisation with local species or human-mediated gene flow (e.g. Bubac & Spellman, 2016; Robalo et al., 2020). This raises the question: is the reed warbler an exception, or is this perhaps more common in nature than has previously been thought?

Apart from the expectation of consecutive founder effects and drift eroding genetic diversity during range expansions (Excoffier et al., 2009), it is also well-known that a few migrants per generation can have a large impact on maintaining diversity (Mills & Allendorf, 1996). Most studies that fail to identify a considerable loss of genetic diversity in association with invasions or environment-driven expansions identify long-distance dispersal (LDD) as a key process maintaining variation at the range front (Berthouly-Salazar et al., 2013; Engler et al., 2016; Song et al., 2013). Can this be the case with the reed warbler? In its range core, the species’ dispersal kernel is characterised by strong philopatry, some short-distance dispersal, and very rare LDD events (Davies, 2019; Hałupka et al., 2022; Paradis et al., 1998). There are estimates from Britain, for example, that short-distance dispersal alone cannot explain the recent rate of reed warblers’ range expansion into Scotland (Davies, 2019). However, considering the rate of expansion and the maintained genetic diversity in Northern Europe, it does not seem plausible that the colonisation of Fennoscandia and the Baltic countries would have been driven by only a few long-distance-dispersing individuals, either. Perhaps the expansion may instead have been accompanied by changes in expansion-facilitating traits, such as increased dispersal propensity, a phenomenon documented in other expanding species (e.g. Duckworth & Badyaev, 2007; Phillips et al., 2006).

An interesting pattern in the data is that apart from the slightly more pronounced genetic structuring among the southernmost sampling sites, levels of differentiation are very similar across the European range, even though the northern populations have only had <150 years to drift apart genetically compared to the thousands of years since postglacial expansion in the Central European range core (Arbabi et al., 2014). This could reflect different processes (or their combinations) behind the observed pattern of isolation by distance, such as isolation by colonisation (IBC), which can create very similar patterns to IBD in the case of serial colonisation events (Orsini et al., 2013). Founder effects can lead to patterns of divergence persisting even for thousands of generations (Boileau et al., 1992), and the differentiation can further be preserved if initial colonisers have ecological or evolutionary advantage over later immigrants (Orsini et al., 2013). Different processes coincidentally creating roughly equal amounts of differentiation at the range core and edge could also explain the ambiguous results of conStruct, where a very small amount of variation was supported as a significant departure from pure IBD in the spatial model, despite all replicates not agreeing on the same split of groups. Also in the PCA, the newly colonised sites did not fully reflect geography and instead formed a cline of differentiation roughly in the order of colonisation, possibly more fitting with IBC. Isolation by environment (IBE) is also a possibility, especially when selective pressures covary with geography within the reed warbler range (Sætre et al., 2022, preprint; Smith et al., 2025, preprint). A recent whole genome sequencing study of a comparably extensive range expansion of Anna’s hummingbird (*Calypte anna*; Adams et al., 2023) reported slight but significant reductions in nucleotide diversity at the range edge but no range-wide population structure, seemingly opposite to the reed warblers with spatial structuring but fully maintained levels of diversity. The authors discuss the possibility that during their timeframe (30-60 years since colonisation of the range edge), allele frequency differences might not have had time to build up. It is possible that the slightly longer time in our case (up to 150 years) has made a difference in pronouncing differentiation and replenishing diversity. However, in terms of generation time (4.3 years for the reed warbler and 2.3 years for Anna’s hummingbird; International Union for Conservation of Nature, 2024), the timescales of the two expansions are very similar and the differences might be explained by other factors. Disentangling the processes behind the observed pattern requires assessment of neutral and adaptive variation separately (Adams et al., 2023; Orsini et al., 2013), making it a promising direction for further research.

### 4.3 Why was the reed warbler so successful in expanding its range?

Apart from some potentially expansion-limiting factors (e.g., low dispersal propensity and habitat specialisation), the reed warbler has had good prerequisites for a successful range expansion, including a large distribution and census population size, high dispersal capabilities, and a possibility for niche-tracking due to newly available habitat (likely a result of eutrophication, decreased grazing pressure, and global warming; Jutila, 2001; Virkkala et al., 2005; von Numers, 2011). These all have been found to be positively associated with range expansion success in previous studies (MacLean & Beissinger, 2017; Mair et al., 2014; Pacifici et al., 2020). The colonising success of the Central and Western European reed warblers might be a consequence of newly available habitat aligning with their inherited migratory bearing (Bensch 1999, Wynn et al. 2022). Migratory overshooting – where individuals extend their migratory journey beyond their typical breeding areas (Newton, 2023) – is particularly likely for juvenile birds, whose navigational inexperience may predispose them to explore new habitats. Such movements may be further reinforced by post-fledging dispersal, a period thought to serve as a mechanism for searching for future breeding areas in juvenile migratory birds, including reed warblers (Nielsen & Bensch, 1995; Mukhin et al. 2005). Once northern habitats are colonised, subsequent generations may integrate magnetic inclination cues from the new range into their navigational toolkit (Wynn et al. 2022), stabilising the population. Another possible mechanism driving the SW-NE expansion could simply be the geographical proximity of SW-migrating populations to the northern range limit, with SE-migrating populations being possibly restricted to areas southeast of Czechia due to greater geographic distances and barriers (e.g. the Carpathian Mountains). The coast of the Baltic Sea also stretches from southwest towards northeast, possibly with more continuous reed bed habitat than what would be available for birds dispersing through continental Europe. The SW-migrating populations would therefore have had both the advantage of shorter distances to newly available habitats and the alignment of their inherited migratory direction with the Baltic coastline, likely providing a natural corridor for dispersal. From a resilience point of view, our results support the importance of suitable habitat networks for successful range expansions, even in the case of highly mobile species.

### 4.4 Conclusions

As anthropogenic environmental change is affecting ecosystems across the globe, it is valuable to understand the consequences of range shifts not only in rare or invasive species, but also for the native and abundant species that make significant contributions to ecosystem functioning (McGeoch & Latombe, 2016). Here we present a rare case where no reductions in overall levels of genetic diversity could be detected despite a rapid and geographically extensive expansion of a philopatric long-distance migrant, the reed warbler. Even with a genome-wide dataset and the use of statistics that are highly sensitive to past founder effects, the only persisting signal could be detected in private alleles while levels of nucleotide diversity and allelic richness were highly consistent across the species’ European breeding range. More theoretical exploration would be beneficial to assess the value of private allelic richness in detecting range expansions in scenarios with different temporal or spatial scales and varying levels of gene flow and admixture. Unlike other measures of genetic diversity, private alleles are always defined in relation to the populations that have been sampled. Testing and interpreting patterns in these alleles therefore highly benefits from previously identified expansion pathways, illustrating the value of combining genomic and ecological data. Reconstructing the expansion history also enables biologically relevant comparisons of genetic and phenotypic changes between the source and range edge populations. How local adaptation persists under considerable gene flow is another key question likely to benefit from integrative approaches combining molecular, ecological and historical data (Rönkä et al., 2024). Importantly, in-depth studies of the origins and genomic consequences of range shifts pave the road for disentangling the mechanisms underlying expansion success, such as possible changes in expansion-facilitating behaviours.

## Acknowledgements

We thank the many people involved in ringing reed warblers and contributing to EURING databases across Europe, and those who assisted in collection of blood samples in the field (including those previously mentioned in Procházka et al. 2011 and Sætre et al. 2022). New samples collected in Finland and Estonia were possible due to assistance from Julius Mäkinen and Hanna Huitu, and advice from Aleksi Lehikoinen, Margus Ellermaa, and Vallo Tilgar. In addition, we thank Kirsi Kähkönen and Airi Lamminmäki for technical assistance in the Molecular Ecology & Systematics lab at the University of Helsinki, Nikita Chernetsov for helpful suggestions on the historical literature, the EvoConGen research group, Petri Kemppainen, and Jochen Wolf for stimulating discussions about the methods and results, and CSC **–** IT Center for Science for computational resources. NB was supported by the Doctoral Programme in Wildlife Biology (LUOVA), University of Helsinki, and the work was funded by Academy of Finland project grant no. 333801 and HiLIFE Fellows grant to RT.

## Ethics statement

All blood samples were collected under permission from animal ethics experimentation boards and with permission to derogate from wildlife protection laws in respective countries. In Finland these were granted by the Project Authorization Board of the Regional State Administrative Agency (ESAVI/7857/2018) and the Centre for Economic Development, Transport and the Environment (VARELY/758/2018), in Estonia permission was granted by Keskkonnaamet (permit number 02-2019). For samples from other locations, see Procházka et al. 2011, Ciloglu et al., 2019, and Sætre et al. 2022.

## Data Accessibility and Benefit-Sharing

### Data accessibility statement

The raw sequence data is available on NCBI SRA (BioProject PRJNA1217894). The filtered datasets, RAD-seq barcodes, and other input files are available from the Dryad Digital Repository: https://doi.org/10.5061/dryad.bcc2fqzrw. All scripts are available from the GitHub repository: https://github.com/norabergman/Reed-warbler-range-expansion. The reference genome for *Acrocephalus scirpaceus* (bAcrSci1) used in this study was published in Sætre et al. 2021 and is available on the European Nucleotide Archive (BioProject PRJEB45715).

### Benefit-sharing statement

Benefits from this research accrue from the sharing of our data and results on public databases as described above.

## Author contributions

**Conceptualization**: Nora M. Bergman, Rose Thorogood, and Katja Rönkä. **Data curation**: Nora M. Bergman, Petteri Lehikoinen, Camilla Lo Cascio Sætre, Frode Fossøy, and Katja Rönkä. **Formal analysis**: Nora M. Bergman, Petteri Lehikoinen, and Katja Rönkä. **Funding acquisition:** Nora M. Bergman, Petteri Lehikoinen, Staffan Bensch, Camilla Lo Cascio Sætre, Fabrice Eroukhmanoff, Petr Procházka, Bård G. Stokke, Rose Thorogood, and Katja Rönkä. **Investigation:** Nora M. Bergman, Petteri Lehikoinen, Edward Kluen, Camilla Lo Cascio Sætre, Fabrice Eroukhmanoff, Rose Thorogood, and Katja Rönkä. **Methodology:** Nora M. Bergman, Petteri Lehikoinen, Edward Kluen, Craig R. Primmer, and Katja Rönkä. **Project administration:** Nora M. Bergman, Fabrice Eroukhmanoff, Rose Thorogood, and Katja Rönkä. **Resources:** Nora M. Bergman, Petteri Lehikoinen, Staffan Bensch, Camilla Lo Cascio Sætre, Fabrice Eroukhmanoff, Frode Fossøy, Petr Procházka, William J. Smith, Bård G. Stokke, Rose Thorogood, and Katja Rönkä. **Software:** Nora M. Bergman and Katja Rönkä. **Supervision:** Fabrice Eroukhmanoff, Craig R. Primmer, Rose Thorogood, and Katja Rönkä. **Validation:** Nora M. Bergman and William J. Smith. **Visualization:** Nora M. Bergman, Petteri Lehikoinen, and Rose Thorogood. **Writing - original draft:** Nora M. Bergman. **Writing - review & editing:** Nora M. Bergman, Petteri Lehikoinen, Edward Kluen, Staffan Bensch, Fabrice Eroukhmanoff, Frode Fossøy, Petr Procházka, William J. Smith, Bård G. Stokke, Craig R. Primmer, Rose Thorogood, and Katja Rönkä.

