## Supplemental Information for "Combined evidence reveals the origin of a rapid range expansion despite retained genetic diversity and a weak founder effect"

### Table of Contents:


| | |
|---|---|
| **Table S1.** | Page 2 |
| **Supplement S2.** | Page 3 |
| **Figure S3.** | Page 4 |
| **Table S4.** | Page 5 |
| **Figure S5.** | Page 6 |
| **Figure S6.** | Page 7 |
| **Table S7.** | Page 8 |
| **Figure S8.** | Page 9 |
| **Figure S9.** | Page 10 |
| **Figure S10.** | Page 11 |
| **Figure S11.** | Page 12 |
| **Figure S12.** | Page 13 |
| **Supplement S13.** | Page 14 |
| **Figure S14.** | Page 15 |
| **Figure S15.** | Page 16 |
| **Figure S16.** | Page 17 |

**Table S1.** Information about the sampling sites and samples included in the final analyses.

| *Country* | *Locality* | *Site code* | *Longitude* | *Latitude* | *Sampling year* | *Sequencing year* | *Sample collector* |
|---|---|---|---|---|---|---|---|
| *Czechia (LU)* | Lužice | CZLU | 17.07 | 48.85 | 2004 | 2017 | P. Procházka |
| *Czechia (OS)* | Osík | CZOS | 16.28 | 49.84 | 2004 | 2017 | P. Procházka |
| *Germany (DG)* | Diergarten | DEDG | 6.10 | 51.23 | 2003 | 2017, 2021 | B. Stokke |
| *Germany (MH)* | Mohrhof | DEMH | 10.85 | 49.66 | 2003 | 2017, 2021 | B. Stokke |
| *Denmark* | Arresø | DK | 12.12 | 55.97 | 2004 | 2021 | B. Stokke |
| *Estonia* | Pärnu | EE | 24.53 | 58.38 | 2019 | 2021 | R. Thorogood / E. Kluen |
| *Spain* | Ebro Delta | ES | 0.79 | 40.73 | 2004 | 2021 | B. Stokke |
| *Finland (E)* | Porvoo | FIE | 25.67 | 60.34 | 2018 | 2021 | R. Thorogood / E. Kluen |
| *Finland (W)* | Kustavi, Taivassalo | FIW | 21.50 | 60.56 | 2018 | 2021 | R. Thorogood / E. Kluen |
| *France* | Trunvel | FR | -4.35 | 47.90 | 2004 | 2017 | B. Stokke |
| *Croatia* | Neretva | HR | 17.45 | 43.05 | 2005 | 2017 | P. Procházka |
| *Italy* | Lago Salso | IT | 15.87 | 41.56 | 2017 | 2017 | F. Eroukhmanoff |
| *Lithuania* | Ventės ragas | LT | 21.24 | 55.35 | 2003 | 2021 | B. Stokke |
| *Norway* | Hemnessjøen | NO | 11.46 | 59.74 | 2002 | 2017 | B. Stokke |
| *Poland* | Milicz | PL | 17.31 | 51.53 | 2003, 2006 | 2021 | B. Stokke |
| *Romania (TR)* | Lacul Ştiucilor | ROTR | 23.90 | 46.97 | 2003 | 2021 | B. Stokke |
| *Romania (DD)* | Lacul Sinoe | RODD | 28.87 | 44.63 | 2004 | 2021 | B. Stokke |
| *Sweden* | Krankesjön | SE | 13.48 | 55.70 | 2016 | 2021 | S. Bensch |
| *Slovakia* | Trnava | SK | 17.58 | 48.37 | 2005 | 2017 | P. Procházka |
| *Turkey* | Mogan Gölü | TR | 32.79 | 39.76 | 2005 | 2017 | P. Procházka |

### S2. Changes to the manufacturer's (DNeasy Blood & Tissue Kit; QIAGEN Inc., California, U.S.A) DNA extraction protocol

Samples stored in Queen's Lysis Buffer (QLB):

- Mixed 125 µl of blood stored in QLB, 75 µl PBS and 20 µl Proteinase K. Mixed thoroughly by vortexing and incubated at 56°C overnight (keeping the mix at 300).
- The next day, added 4 µl RNase A (100 mg/ml), mixed by vortexing and incubated 2 min at room temperature. Then followed manufacturer's protocol (with changes mentioned below) from step 2.

Samples stored in ethanol:

- Separated an approximately 1x1 mm piece of the sample blood pellet, as solid as possible, into a prepared tube.
- To evaporate all ethanol, heated the tubes with the lids open on a heat block for 1 hour at 56°C, all tubes loosely covered in foil and a cleaned heat block lid.
- Added 180 µl Buffer ATL and 20 µl Proteinase K in each tube. Mixed thoroughly by vortexing.
- Incubated at 56°C overnight (keeping the mix at 300). Vortexed shortly (~3 s) a few times during incubation to help dissolve the pellets.
- The next day, vortexed all samples (~3 s), added 4 µl RNase A (100 mg/ml), mixed by vortexing and incubated 2 min at room temperature. Then followed manufacturer's protocol (with changes mentioned below) from step 2.

Samples stored on Whatman® FTA® cards:

- Using a Harris Uni-Core™ punch tool (Ø: 1.2 mm) and a Harris cutting mat, punched 3 holes in the sample on each FTA card and placed the cut-out circles in prepared tubes. The equipment were sterilised between samples.
- Added 180 µl Buffer ATL and 20 µl Proteinase K in each tube. Mixed thoroughly by vortexing.
- Incubated at 56°C overnight (keeping the mix at 300).
- The next day, vortexed all samples (~3 s), added 4 µl RNase A (100 mg/ml), mixed by vortexing and incubated 2 min at room temperature. Then followed manufacturer's protocol (with changes mentioned below) from step 2.

Changes to the later steps of the protocol (for all samples):

- In steps 5 & 6, incubated 5 min at room temperature after pipetting the wash buffer into the spin columns, which improved extract purity.
- In step 7, eluted the DNA into 100 µl buffer EB (QIAGEN Inc., California, U.S.A) and incubated at room temperature for 5 mins before centrifuging. Finally, prepared a second eluate into 60 µl buffer EB.

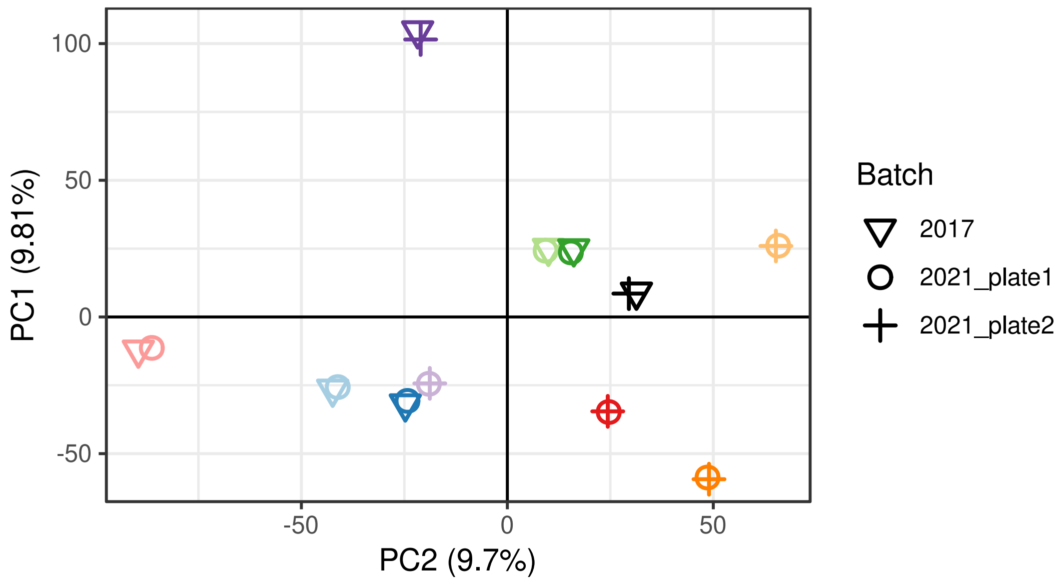


**Figure S3.** The principal component analysis (PCA) of the control samples, each sequenced in two out of three sequencing batches. The shape of the point indicates the sequencing batch, and the colour indicates the individual ID (each individual sequenced twice across batches). The samples cluster by sample identity and not by batch, suggesting the robustness of population structure analyses against any batch differences.

**Table S4.** Results of the between-batch comparisons from paired, two-tailed t-tests. Samples were sequenced in three batches: 2017, 2021_plate1, and 2021_plate2. All comparisons between 2021_plate1 and 2021_plate2 are below the statistical significance level ($p > 0.05$), while all comparisons between the combined 2021 plates and the 2017 batch show slight but statistically significant differences.

| **2021_plate1 vs. 2021_plate2** | **2021 (both plates combined) vs. 2017** |
|---|---|
| Mean depth per individual:<br>t = 0.88216, df = 3, p = 0.44<br>Mean difference: 6.854 | Mean depth per individual:<br>t = -2.7237, df = 6, **p = 0.03**<br>Mean difference: -18.149 |
| Missigness per individual:<br>t = 0.86142, df = 3, p = 0.45<br>Mean difference: 0.003 | Missigness per individual:<br>t = -5.0632, df = 6, **p = 0.002**<br>Mean difference: -0.038 |
| Mean heterozygosity per individual:<br>t = -2.2868, df = 3, p = 0.11<br>Mean difference: -0.003 | Mean heterozygosity per individual:<br>t = 6.367, df = 6, **p < 0.001**<br>Mean difference: 0.037 |
| Expected heterozygosity per individual:<br>t = -0.2708, df = 3, p = 0.80<br>Mean difference: -0.0001 | Expected heterozygosity per individual:<br>t = 2.8631, df = 6, **p = 0.03**<br>Mean difference: 0.001 |

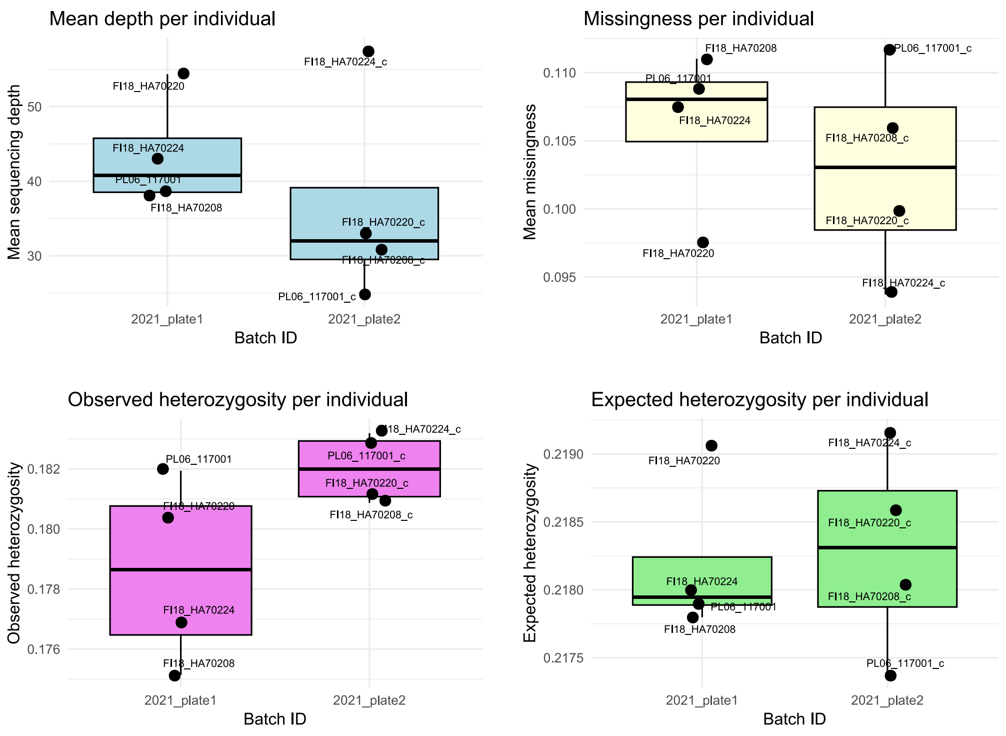


**Figure S5.** Box plots showing the individual-based statistics (mean depth of coverage, mean proportion of missing data, observed proportion of heterozygous sites, and expected proportion of heterozygous sites) in the sequencing batches 2021_plate1 and 2021_plate2.

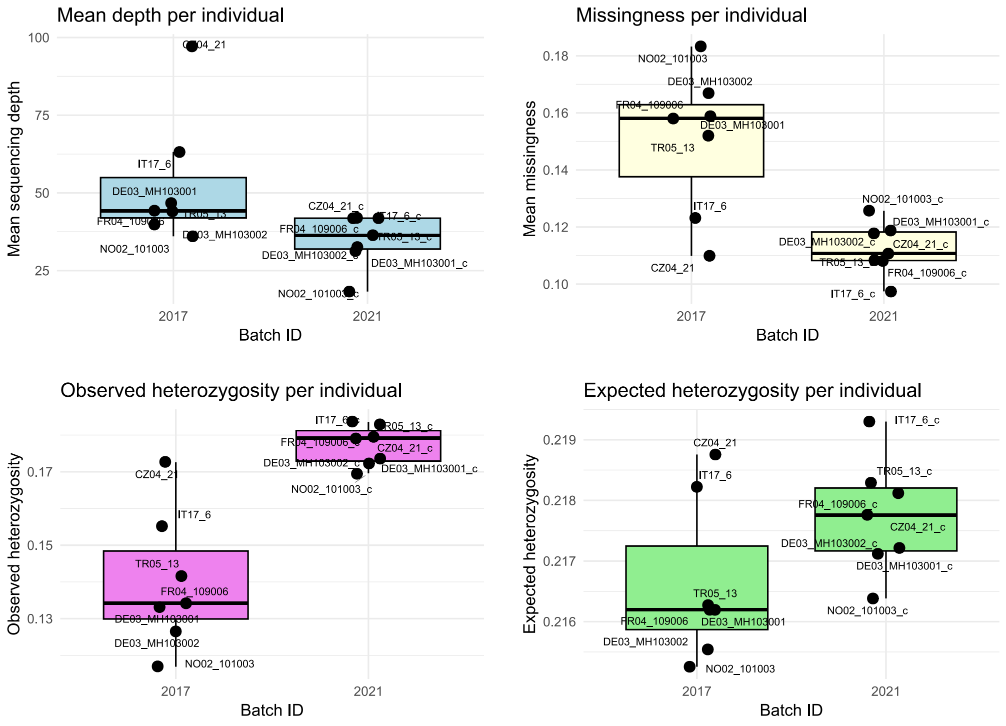


**Figure S6.** Box plots showing the individual-based statistics (mean depth of coverage, mean proportion of missing data, observed proportion of heterozygous sites, and expected proportion of heterozygous sites) in the combined 2021 batches (2021_plate1 & 2021_plate2) and the 2017 batch.

**Table S7.** The filtered datasets that were used in the final analyses.

| Dataset | Description | Analyses | N (ind.) | Sampling sites | #Loci; #SNPs |
|---|---|---|---|---|---|
| **1) Full, linkage-pruned SNP dataset** | All sampling sites included; SNPs pruned by physical distance | PCA, conStruct, $F_{ST}$ | 171 | 20 | 35564; 35564 |
| **2) Full, linkage-pruned RAD locus dataset** | All sampling sites included; full RAD loci pruned by physical distance | fineRADstructure | 171 | 20 | 35463; 246429 |
| **3) Batch-effect-free, linkage-pruned SNP dataset** | Only sampling sites from 2021 batches; SNPs pruned by physical distance | RangeExpansion | 102 | 12 | 34547; 34547 |
| **4) Batch-effect-free, full RAD locus dataset (not linkage-pruned)** | Only sampling sites from 2021 batches; not linkage-pruned | $\pi$, (private) allelic richness | 102 | 12 | 78164; 246721 |

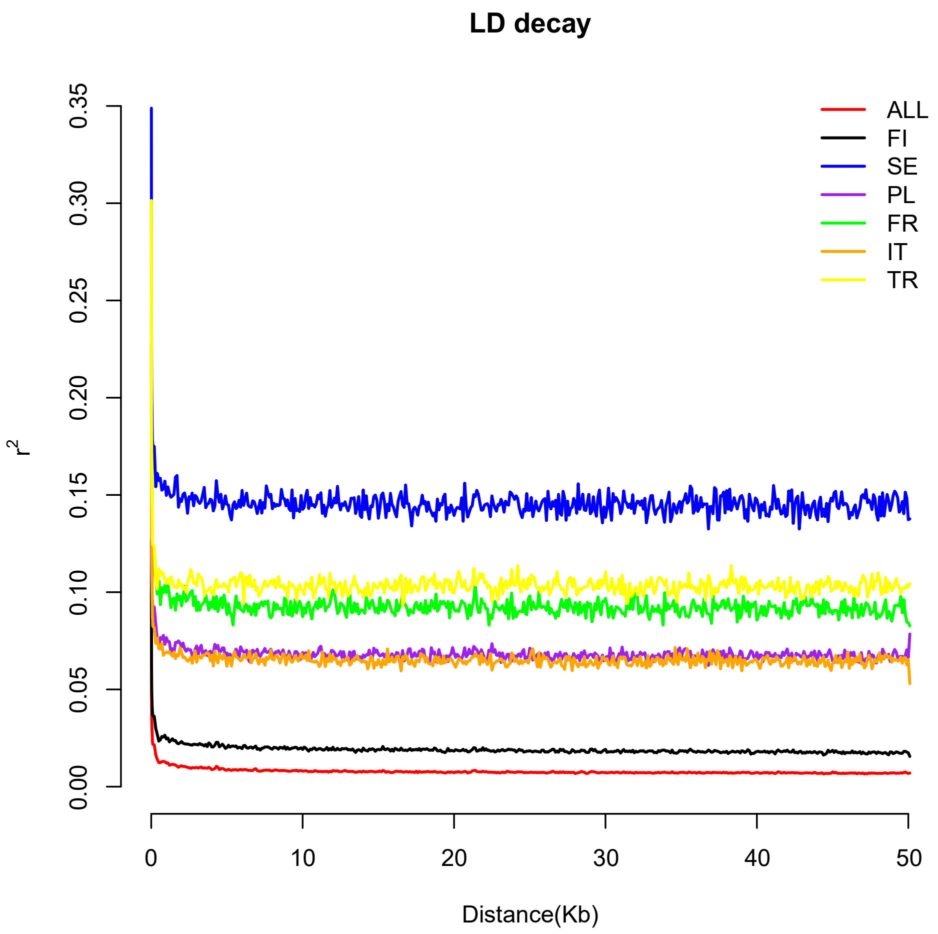


**Figure S8.** The linkage disequilibrium decay distances calculated with PopLDdecay (v3.42; Zhang et al., 2019). Lines with different colours indicate the values of LD ($r^2$) over physical distance (Kb) in different sampling sites before downsampling to a maximum of 10 individuals per site. The red line shows the LD decay distance across all samples. The differences in the different plateauing values between lines are due to the number of samples from the corresponding sampling site (in this plot, i.e. before downsampling, $n_{SE}$ = 6, $n_{TR}$ = 8, $n_{FR}$ = 9, $n_{IT}$ = 13, $n_{PL}$ = 13, $n_{FI}$ = 84, $n_{ALL}$ = 260). However, the decay pattern is similar across all tested sampling sites and plateaus around 3 Kb, which we set as the filtering threshold for analyses that assume unlinked markers.

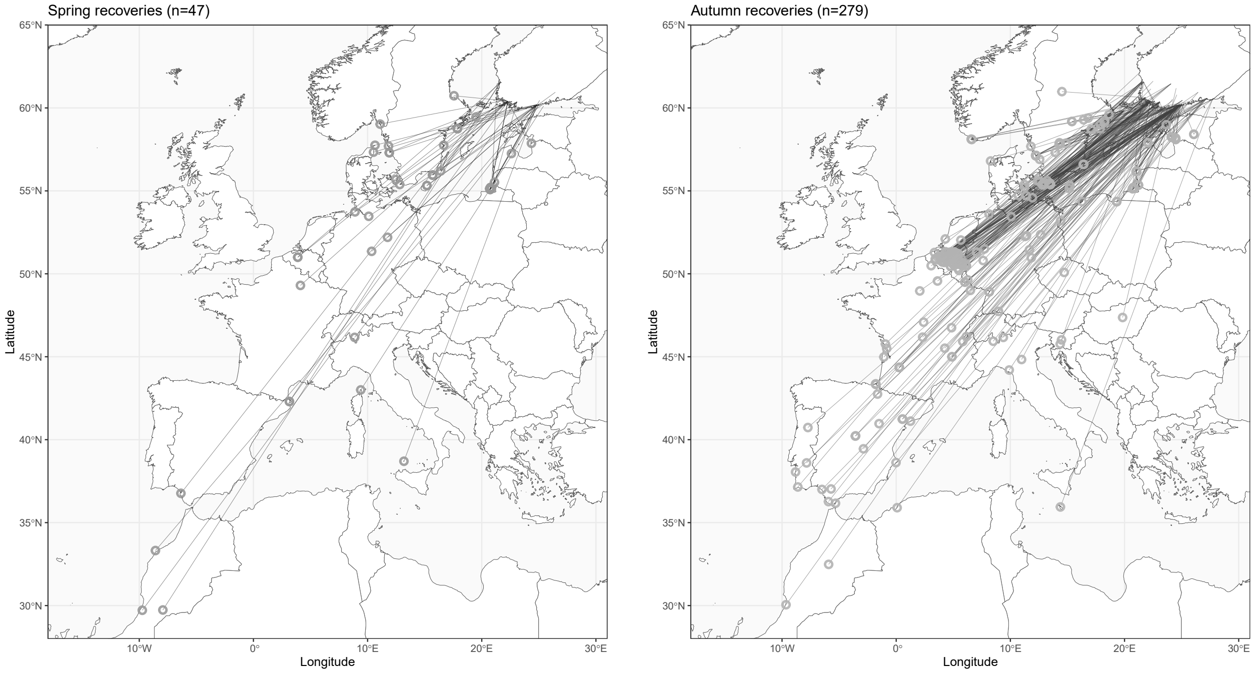


**Figure S9.** The re-encounters of reed warblers ringed in Finland and captured on migration in other countries (years 1969-2023), divided into spring recoveries (left; January–June) and autumn recoveries (right; July–December).

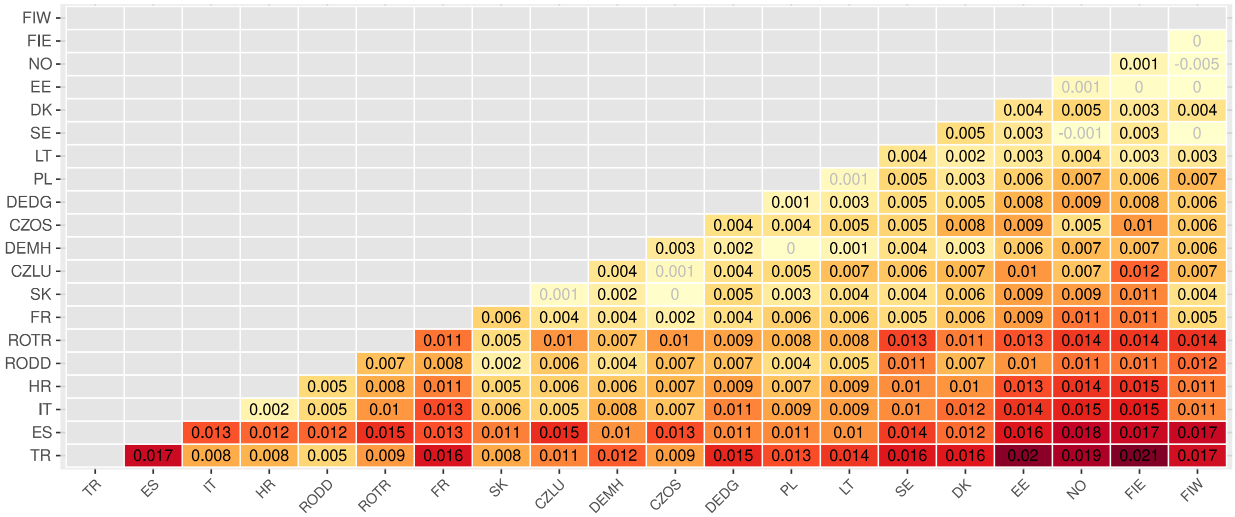


**Figure S10.** Pairwise $F_{ST}$ between the sampling sites, ordered by latitude. Darker colours indicate higher differentiation. Significant $F_{ST}$ values ($p < 0.05$) are reported in black text, and non-significant values in light grey.

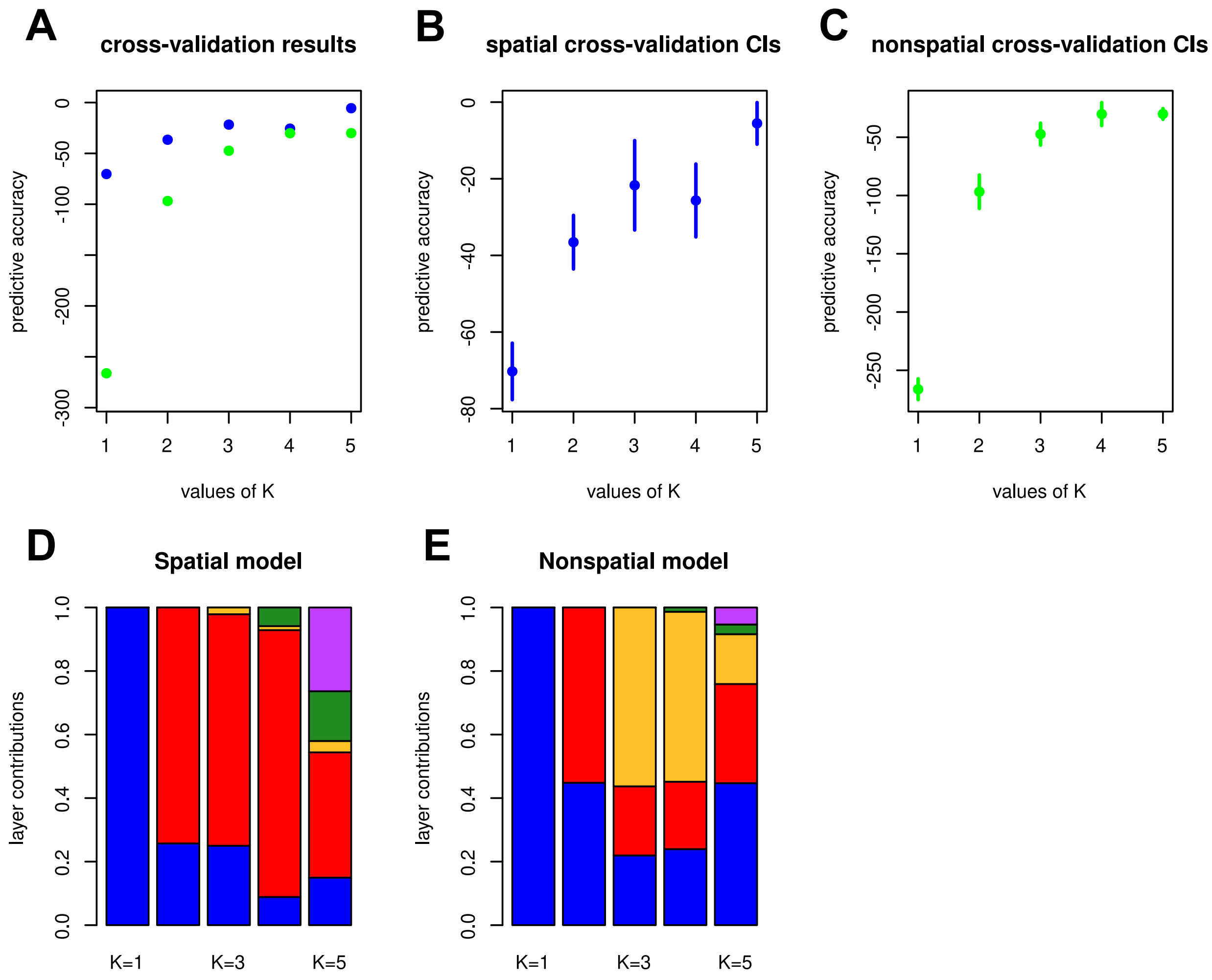


**Figure S11. A.** Cross-validation showing the predictive accuracy of the spatial model (blue) and non-spatial model (green) across K = 1-5. The spatial model has a higher predictive accuracy at all values of K. **B.** The cross-validation values with confidence intervals for the spatial model. **C.** The cross-validation values with confidence intervals for the nonspatial model. **D.** Layer contributions of the spatial model to the total covariance of the model at K = 1-5. **E.** Layer contributions of the nonspatial model to the total covariance of the model at K = 1-5.

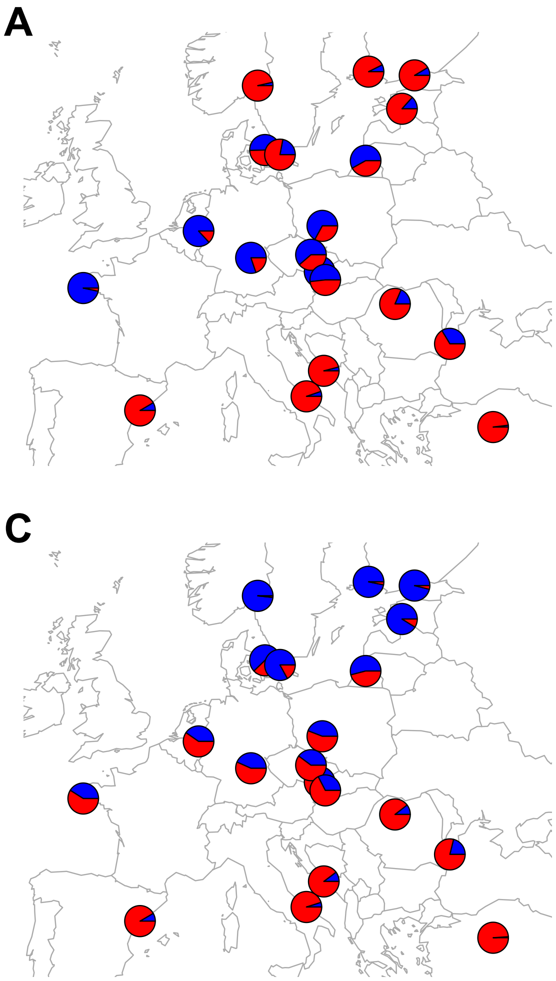


**Figure S12.** The four replicates of the best-supported model (spatial model with K = 2) did not converge on the same split of layers, likely due to the small value of the “shared drift” parameter phi within each layer and therefore the small amount of variation separating the layers (phi(layer1) = 1.34e-06 – 1.63e-05 per replicate, phi(layer2) = 4.62e-06 – 4.83e-05 per replicate). Two replicates (A-B) suggest that the northern and southern samples would share more ancestry from the same layer than expected under IBD, one replicate (C) splits the northern and southern samples into separate layers, and the last replicate (D) also is roughly similar to A and B but with somewhat different ancestry proportions and Spain now grouping with the Central European layer. The discrepancies between the replicates with K = 2 may indicate that the most biologically relevant grouping could actually be a spatial model with K = 1.

**S13.** Genetic diversity results (nucleotide diversity, π & allelic richness, AR)

Genetic diversity values and t-test grouping (sampling sites arranged by latitude from north to south).

| Sampling site | Nucleotide diversity (π) | Allelic richness (standardised to G = 10) | T-test grouping |
|---|---|---|---|
| FIW | 0.00595 | 1.47020 | EDGE |
| FIE | 0.00604 | 1.47689 | EDGE |
| EE | 0.00607 | 1.47870 | EDGE |
| DK | 0.00605 | 1.47903 | EDGE |
| SE | 0.00586 | 1.47038 | EDGE |
| LT | 0.00606 | 1.48007 | EDGE |
| PL | 0.00608 | 1.48102 | CORE |
| DEDG | 0.00601 | 1.47337 | CORE |
| DEMH | 0.00598 | 1.47909 | CORE |
| ROTR | 0.00597 | 1.47202 | CORE |
| RODD | 0.00602 | 1.47809 | CORE |
| ES | 0.00598 | 1.46919 | CORE |

Simple linear regressions:

Nucleotide diversity π ~ Latitude: $\beta = 9.787e\text{-}07$, $SE = 3.133e\text{-}06$, $t = 0.312$, $p = 0.761$

Allelic richness ~ Latitude: $\beta = 0.0001294$, $SE = 0.0002143$, $t = 0.604$, $p = 0.559$

Two-tailed t-tests (EDGE vs. CORE):

Nucleotide diversity π: $t = 0.044082$, $df = 7.2794$, $p = 0.966$
Mean π in the recently colonised range, Fennoscandia & Baltic countries (EDGE): 0.006005
Mean π in the other sites (CORE): 0.006007

Allelic richness: $t = -0.15857$, $df = 9.9865$, $p = 0.877$
Mean AR in the recently colonised range, Fennoscandia & Baltic countries (EDGE): 1.475878
Mean AR in the other sites (CORE): 1.475463

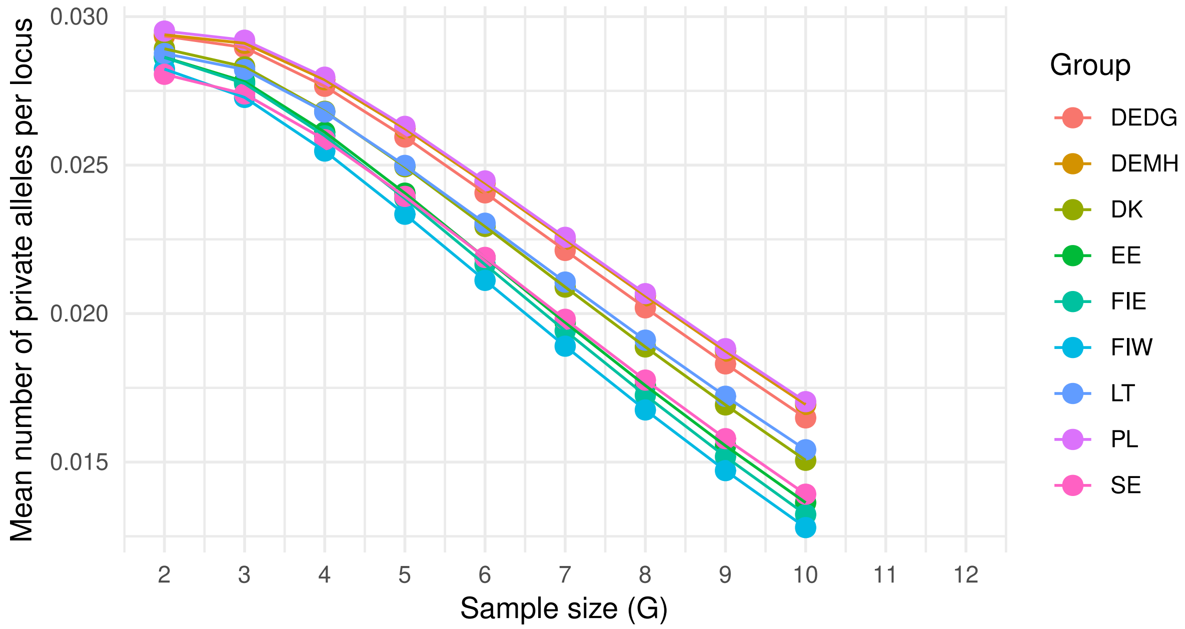


**Figure S14.** The expected private allelic richness for different standardised sample sizes (G denotes the number of allele copies per sampling site). While the number of alleles is not saturated at G = 10 (the lowest sample size across the sampling sites in our dataset), the relative differences between sampling sites seem to persist and become even more pronounced with increasing sample size.

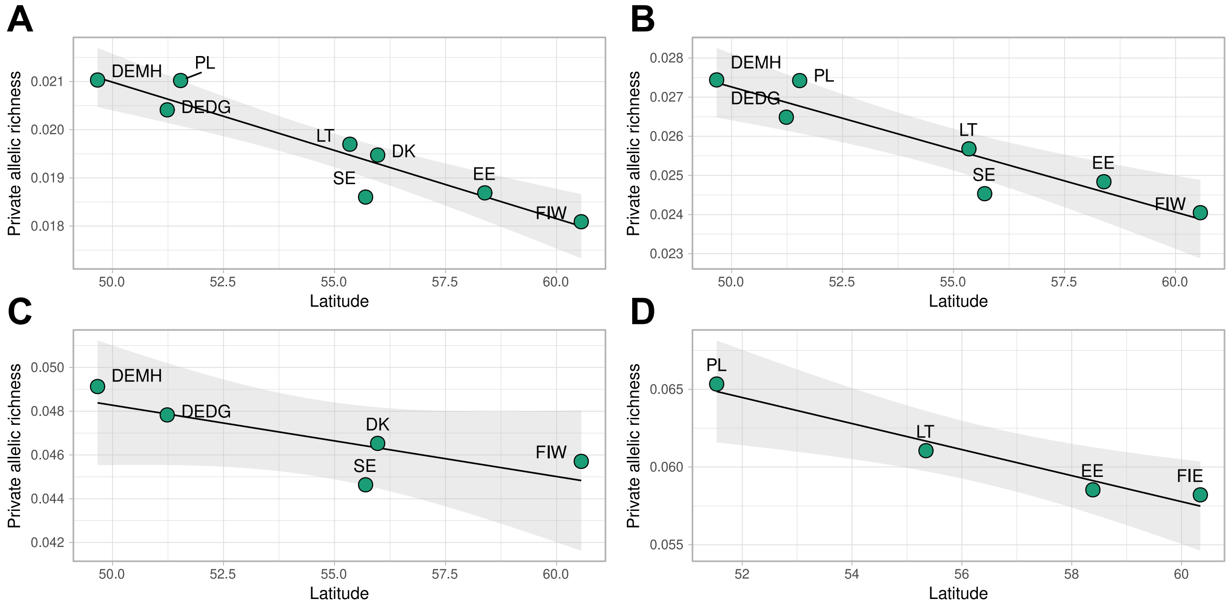


**Figure S15.** The pattern of private allelic richness persists when excluding sites in close geographical proximity with each other (A-B) or looking at both sides of the Baltic Sea separately (C-D). Standardised sample size G = 10.

**A.** Sampling sites included: DEMH, DEDG, PL, LT, SE, DK, EE, FIW (excluding one of the two Finnish sites FIE). Private allelic richness ~ Latitude: β = -2.834e-04, SE = 4.045e-05, t = -7.006, p = 0.000421, $R^2$ = 0.8911

**B.** Sampling sites included: DEMH, DEDG, PL, LT, SE, EE, FIW (excluding one of the two Finnish sites FIE, and Denmark DK due to geographical proximity with Sweden). Private allelic richness ~ Latitude: β = -3.200e-04, SE = 5.575e-05, t = -5.74, p = 0.00225, $R^2$ = 0.8682

**C.** Sampling sites included: DEMH, DEDG, DK, SE, FIW (sites along the western coast of the Baltic Sea). Private allelic richness ~ Latitude: β = -0.0003259, SE = 0.0001425, t = -2.287, p = 0.1063, $R^2$ = 0.6355

**D.** Sampling sites included: PL, LT, EE, FIE (sites along the eastern coast of the Baltic Sea). Private allelic richness ~ Latitude: β = -0.0008376, SE = 0.0001294, t = -6.473, p = 0.02304, $R^2$ = 0.9544

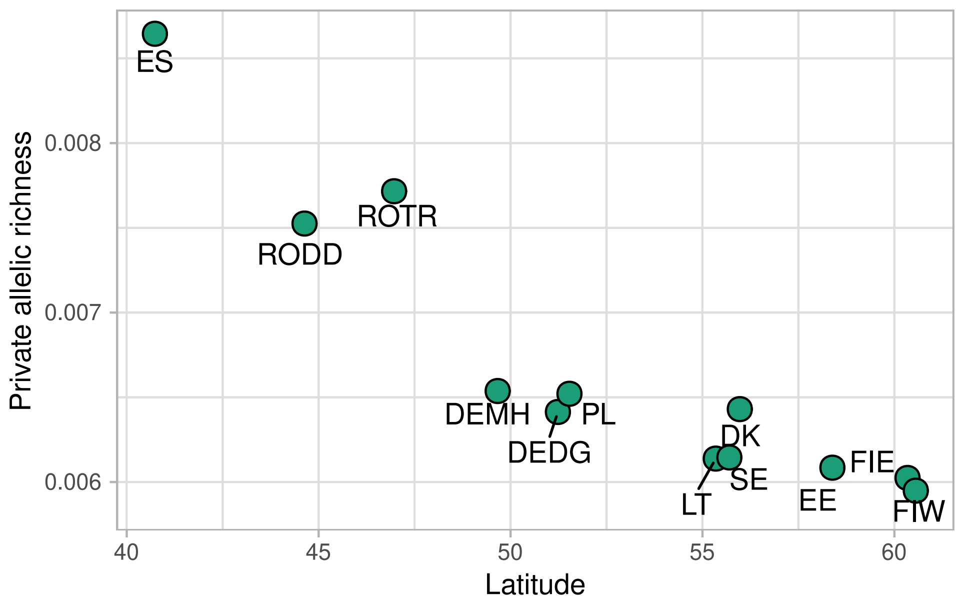


**Figure S16.** Private allelic richness calculated across all sampling sites in the data set. Standardised sample size G = 10.
